# Limitation of phosphate assimilation maintains cytoplasmic magnesium homeostasis

**DOI:** 10.1101/2020.09.05.284430

**Authors:** Roberto E. Bruna, Christopher G. Kendra, Eduardo A. Groisman, Mauricio H. Pontes

**Affiliations:** Department of Pathology and Laboratory Medicine, Penn State College of Medicine, 500 University Drive, P.O. Box 17033, Hershey, PA, 17033, USA; Department of Microbiology and Immunology, Penn State College of Medicine, 500 University Drive, P.O. Box 17033, Hershey, PA, 17033, USA; Department of Microbial Pathogenesis, Yale School of Medicine, 295 Congress Avenue, New Haven, CT 06536, USA; Yale Microbial Sciences Institute, P.O. Box 27389, West Haven, CT, 06516, USA

**Keywords:** Phosphate, ATP, Magnesium, MgtC, *Salmonella*

## Abstract

Phosphorus (P) is an essential component of several core biological molecules. In bacteria, P is mainly acquired as inorganic orthophosphate (Pi). Once in the cytoplasm, Pi is incorporated into adenosine triphosphate (ATP), which exists primarily as a Mg^2+^ salt. Notably, whereas P is essential, excess of cytosolic Pi hinders growth. Here we demonstrate that cytotoxic effects of excessive Pi uptake result from its assimilation into ATP and subsequent disruption of Mg^2+^ dependent processes. We show that *Salmonella enterica* cells experiencing cytoplasmic Mg^2+^ starvation restrict Pi uptake, thereby limiting the availability of an ATP precursor. This response prevents excessive ATP synthesis, overproduction of ribosomal RNA, chelation of free cytoplasmic Mg^2+^ and the destabilization of Mg^2+^-dependent core processes that ultimately hinder bacterial growth and leads to loss of cellular viability. We demonstrate that, even when cytoplasmic Mg^2+^ is not limiting, excessive Pi uptake leads to increased ATP synthesis, depletion of free cytoplasmic Mg^2+^, inhibition of translation and growth. Our results establish that bacteria must restrict Pi uptake to prevent the depletion of cytoplasmic Mg^2+^. Furthermore, they provide a framework to understand the molecular basis of Pi cytotoxicity and reveal a regulatory logic employed by bacterial cells to control P assimilation.

**Importance:** Phosphorus (P) is essential for life. As the fifth most abundant element in living cells, P is required for the synthesis of an array of biological molecules including (d)NTPs, nucleic acids and membranes. Organisms typically acquire environmental P as inorganic phosphate. While essential for growth and viability, excessive intracellular Pi is toxic for both bacteria and eukaryotes. Using the bacterium *Salmonella enterica* as a model, we demonstrate that Pi cytotoxicity is manifested following its assimilation into ATP, which acts as a chelating agent for intracellular cations, most notably, Mg^2+^. These results identify physiological processes disrupted by excessive Pi and elucidate a regulatory logic employed by bacteria to prevent uncontrollable P assimilation.

## Introduction

Phosphorus (P) is an intrinsic component of a large number of biological molecules, including membrane lipids, nucleotides and nucleic acids. This element is required for many central biological functions, including (1) the formation of cellular boundaries, (2) the storage and transfer of chemical energy, (3) the integration and propagation of information in signal transduction pathways, and (4) the storage, transmission and expression of genetic information. Bacterial cells mainly uptake P as inorganic phosphate (PO_4_^-3^; Pi), which is then assimilated in the cytoplasm via its incorporation into adenosine triphosphate (ATP). ATP functions as the main cellular P-carrier molecule, mediating both the transfer of Pi among biological molecules and the release of chemical energy to power energy-dependent processes (1).

Interestingly, while the assimilation of P is essential, excessive cytoplasmic Pi is toxic (2–7). This implies that cells must tightly regulate Pi acquisition and utilization to avoid self-poisoning. Here, we elucidate how bacterial cells coordinate Pi acquisition and consumption to prevent deleterious effects of unbalanced Pi metabolism.

Following assimilation, negative charges from Pi groups in biomolecules are neutralized by positively charged ionic species present in the cytoplasm. As such, the majority of cytoplasmic ATP exists as a salt with positively charged magnesium (Mg^2+^), the most abundant divalent cation in living cells. Indeed, this ATP:Mg^2+^ salt, rather than the ATP anion, functions as the substrate for most ATP-dependent enzymatic reactions (8–10). In enteric bacteria, ATP stimulates the transcription of ribosomal RNAs (rRNA) (11, 12). The synthesis and activity of ribosomes consume the majority of the ATP in the cell (13). During ribosome biogenesis, negative charges from Pi groups in the rRNA backbone chelate large amounts of Mg^2+^ ions. This process reduces electrostatic repulsion among Pi groups in the rRNA backbone, enabling the folding and assembly of functional ribosomes. Not accidentally, ATP and rRNA constitute the largest cytoplasmic reservoirs of Pi *and* Mg^2+^ (9, 11, 13–18). Given this inherent connection between Pi and Mg^2+^, we wondered if cytotoxic effects of excessive Pi uptake result from its assimilation into ATP and subsequent disruption of Mg^2+^ dependent processes in the cytoplasm.

In the Gram negative bacterial pathogen *Salmonella enterica,* prolonged growth in media containing limiting Mg^2+^ induces cytoplasmic Mg^2+^ starvation. This stress promotes the expression of the MgtA, MgtB and MgtC membrane proteins (19–22). MgtA and MgtB function as high-affinity, ATP-dependent Mg^2+^ importers, increasing the Mg^2+^ concentration in the cytoplasm (23, 24). By contrast, MgtC decreases intracellular ATP levels (25), thereby reducing rRNA synthesis, lowering steady-state levels of ribosomes, and slowing translation rates (26). This response reduces levels of assimilated P and, consequently, the quantity of Mg^2+^ required as a counter-ion, effectively preventing the depletion of cytoplasmic Mg^2+^ that is required for the stabilization of existing ribosomes and the maintenance of other vital Mg^2+^-dependent cellular processes (19, 26, 27). Interestingly, MgtC is also expressed during replication of *Salmonella* inside mammalian macrophages, where it promotes bacterial survival and enables the establishment of systemic infections (28, 29). In macrophages, MgtC inhibits the activity of *Salmonella’s* F_1_F_o_ ATP synthase, thereby hindering ATP production via oxidative phosphorylation (30, 31). Yet, mutations in single amino acid residues of the MgtC protein abolish intramacrophage survival of *Salmonella* without affecting its growth and viability during cytoplasmic Mg^2+^ starvation. This indicates that MgtC operates by distinct mechanisms in these two growth conditions (32).

In this paper, we reveal that the limitation of phosphate uptake is essential to maintain cytoplasmic Mg^2+^ homeostasis. We establish that during cytoplasmic Mg^2+^ starvation, MgtC lowers ATP levels by inhibiting Pi uptake, thus limiting an ATP precursor instead of interfering with its enzymatic generation. We demonstrate that, counterintuitively, limitation of exogenous Pi availability rescues translation, promotes growth and restores viability to an *mgtC* mutant. We provide genetic and functional evidence that MgtC hinders the activity of an uncharacterized transporter, which functions as the main Pi uptake system in *Salmonella* and likely other bacterial species. Finally, we establish that even at physiological levels of cytoplasmic Mg^2+^, Pi exerts its toxicity following its incorporation into ATP and subsequent disruption of Mg^2+^-dependent processes. While providing a conceptual framework to understand the underlying basis of Pi cytotoxicity, a phenomenon observed in both bacteria and eukaryotes (3–7, 33–36), our results uncover a regulatory logic employed by bacterial cells for the global control of P assimilation.

## Results

### An F_1_F_o_ synthase-independent mechanism limits ATP accumulation during low cytoplasmic Mg^2+^ stress

In living cells, ATP exists as a Mg^2+^ salt (10, 13). When cells are faced with limiting cytoplasmic Mg^2+^ concentrations, they reduce ATP levels to free Mg^2+^ ions that are required for other cellular processes, such as the assembly of ribosomes (26, 27). In *Salmonella*, this reduction in ATP levels is accomplished by the MgtC membrane protein, which is expressed in response to a number of physiological signals generated by cytoplasmic Mg^2+^ starvation (19, 21, 25, 37, 38). MgtC is also expressed during *Salmonella* replication in mammalian macrophages, where it promotes bacterial survival (28, 29). In macrophages, MgtC functions by inhibiting the activity of *Salmonella’s* F_1_F_O_ ATP synthase, the enzyme responsible for ATP synthesis via oxidative phosphorylation (30). Notably, a *Salmonella mgtC atpB* double mutant strain—containing a genetically inactive ATP synthase—was also reported to harbor lower ATP levels than an *mgtC* single mutant strain during cytoplasmic Mg^2+^ starvation (30). This result led to the notion that MgtC also prevents a non-physiological rise in ATP levels by inhibiting the F_1_F_O_ ATP synthase when *Salmonella* experiences cytoplasmic Mg^2+^ starvation (30, 39, 40). However, because the biochemical function of MgtC during cytoplasmic Mg^2+^ starvation can be genetically separated from its function during intramacrophage replication (32), we sought to reexamine the interpretation of the aforementioned experimental results.

The F_1_F_O_ ATP synthase uses the proton motive force, generated by the respiratory electron transport chain, to synthesize ATP from ADP and Pi (41, 42). Consequently, an *mgtC atpB* double mutant (or any other strain lacking a functional ATP synthase) relies exclusively on fermentative pathways to produce ATP via substrate-level phosphorylation (43). Interestingly, the aforementioned lower ATP levels observed in a *Salmonella mgtC atpB* double mutant were obtained during growth on medium containing glycerol as the carbon source (30). Given that the fermentation of glycerol is extremely inefficient (44, 45), we reasoned that lower ATP levels in the *mgtC atpB* double mutant could simply reflect an inability of this strain to efficiently ferment glycerol. To test this notion, we compared ATP levels in wild-type, *mgtC*, *atpB* and *mgtC atpB Salmonella* strains grown in minimal medium containing readily fermentable glucose as the carbon source and low (10 μM) Mg^2+^, to induce cytoplasmic Mg^2+^ starvation (20, 21). Consistent with our hypothesis, we established that after 5 h of growth, when cytoplasmic Mg^2+^ becomes limiting (26), *mgtC* and *mgtC atpB* strains had 42 and 29-fold higher ATP levels relative to their *mgtC*^+^ isogenic counterparts, respectively (Fig. 1*A*). Hence, a second mutation in *atpB* does not abrogate the intracellular ATP accumulation in an *mgtC*^-^ background provided cells are fed with glucose as the carbon source.

**Figure 1.**
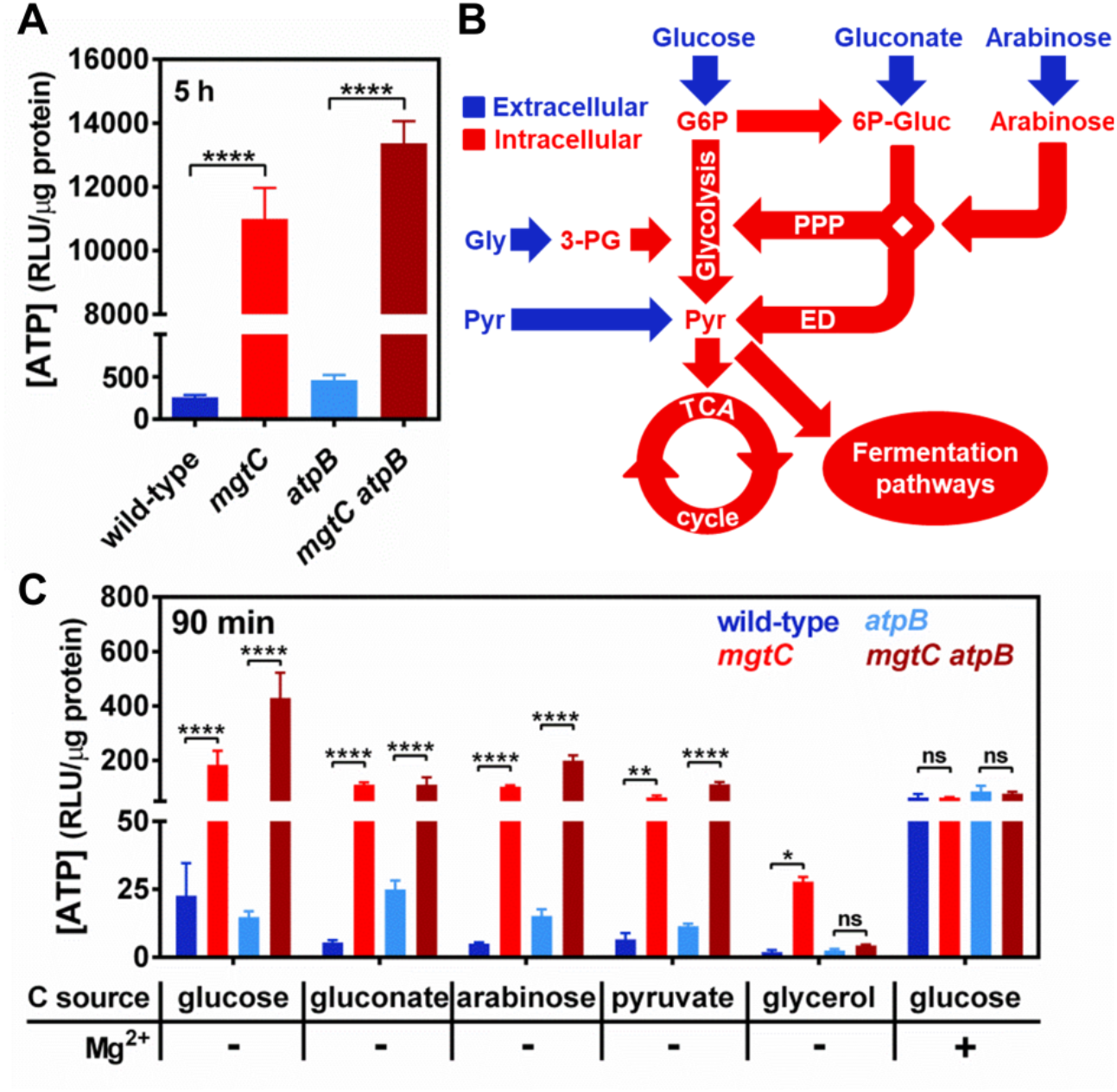
F_1_F_O_ synthase-independent ATP accumulation in an *mgtC* mutant during cytoplasmic Mg^2+^ starvation. *(A)* Intracellular ATP levels in wild-type (14028s), *mgtC* (EL4), *atpB* (MP24) or *mgtC atpB* (MP25) *Salmonella* following 5 h of growth. Cells were grown in MOPS medium containing 10 μM MgCl_2_ and 2 mM K_2_HPO_4_. *(B)* Schematic representation of carbon flow through the central metabolic pathways in bacteria with key extracellular (blue) and intracellular (red) metabolites. G6P, glucose-6-phosphate; 6P-Gluc, 6-Phosphogluconate; Gly, glycerol; Pyr, pyruvate; PPP, pentose phosphate pathway; ED, Entner–Doudoroff pathway. *(C)* Intracellular ATP levels in wild-type (14028s), *mgtC* (EL4), *atpB* (MP24) or *mgtC atpB* (MP25) *Salmonella.* Cells were grown in MOPS medium containing 10 mM MgCl_2_, 2 mM K_2_HPO_4_ and 25 mM glucose until OD_600_ ≈ 0.4, washed thrice, and grown for additional 90 min in MOPS medium supplemented with the indicated carbon source (25 mM glucose, 25 mM sodium gluconate, 30 mM L-arabinose, 50 mM sodium pyruvate, or 50 mM glycerol), and containing 2 mM K_2_HPO_4_ and either 0 (-) or 10 (+) mM MgCl_2_. Means ± SDs of three independent experiments are shown. *P < 0.05, **P < 0.01, ****P < 0.0001, and ns, no significant difference. Two-tailed *t* test *(A);* two-way analysis of variance (ANOVA) with Tukey correction *(C)*.

To further test our hypothesis, we compared ATP levels in wild-type, *mgtC*, *atpB* and *mgtC atpB* strains at 90 min following a nutritional downshift from medium containing high (10 mM) Mg^2+^ and glucose, to media lacking Mg^2+^ and containing one of various carbon sources (Fig. 1*B*). In agreement with our hypothesis, there was a predictable trend in ATP levels among these strains. That is, we determined that during cytoplasmic Mg^2+^ starvation the *mgtC* strain had high ATP levels regardless of the carbon source (Fig. 1*C*). By contrast, the *mgtC atpB* strain had increased ATP levels during growth on readily fermentable carbon sources (glucose, gluconate, arabinose or pyruvate), but was unable to do so during growth on inefficiently fermentable glycerol (Fig. 1*B-C*). As a control, all the strains tested displayed similar ATP levels when resuspended in high Mg^2+^, glucose-containing medium (Fig. 1*C*). Taken together, these results indicate that the low ATP levels previously observed in an *mgtC atpB* strain experiencing cytoplasmic Mg^2+^ starvation (30) are caused by an inability to efficiently ferment glycerol, as opposed to an inability to inhibit the F_1_F_O_ ATP synthase. Furthermore, these results indicate that MgtC inhibits multiple ATP-generating reactions in the cell, not only that one which is carried out by the F_1_F_O_ complex.

### MgtC inhibits inorganic phosphate acquisition and *ipso facto* limits ATP synthesis during cytoplasmic Mg^2+^ starvation

ATP can be synthesized by several catabolic reactions in the cell (46–48). How then can MgtC control the activity of multiple ATP-producing enzymatic reactions? We reasoned that regardless of the identity of the enzymes catalyzing ATP formation, the overall rate of ATP synthesis in the cell could be restricted by the availability of substrates. MgtC could, therefore, function by inhibiting the synthesis or acquisition of an ATP precursor. Interestingly, at the onset of cytoplasmic Mg^2+^ starvation, a temporary shortage in the levels of free cytoplasmic Mg^2+^ destabilizes the bacterial ribosomal subunits (26). The resulting decrease in translation efficiency reduces ATP consumption and, consequently, the recycling of Pi from ATP. This lowers the concentration of cytoplasmic Pi, transiently activating the cytoplasmic Pi-starvation sensing PhoB/PhoR two-component system in *Salmonella* (15). Three lines of evidence led us to hypothesize that MgtC prevents ATP synthesis by limiting Pi influx into the cell. First, PhoB/PhoR activation is hampered in an *mgtC* mutant, indicating that this strain experiences excess cytoplasmic Pi (15). Second, *mgtC* and *mgtC atpB* strains have higher intracellular steady-state Pi levels than wild-type and *atpB Salmonella* strains (Fig. 2*A*), confirming that MgtC prevents the accumulation of intracellular Pi in an *atpB*-independent fashion. Third, when Mg^2+^ and Pi are abundant, ectopic expression of MgtC from a plasmid induces PhoB/PhoR activation, further suggesting that MgtC causes a shortage in cytoplasmic Pi (Fig. S1) (15).

**Figure 2.**
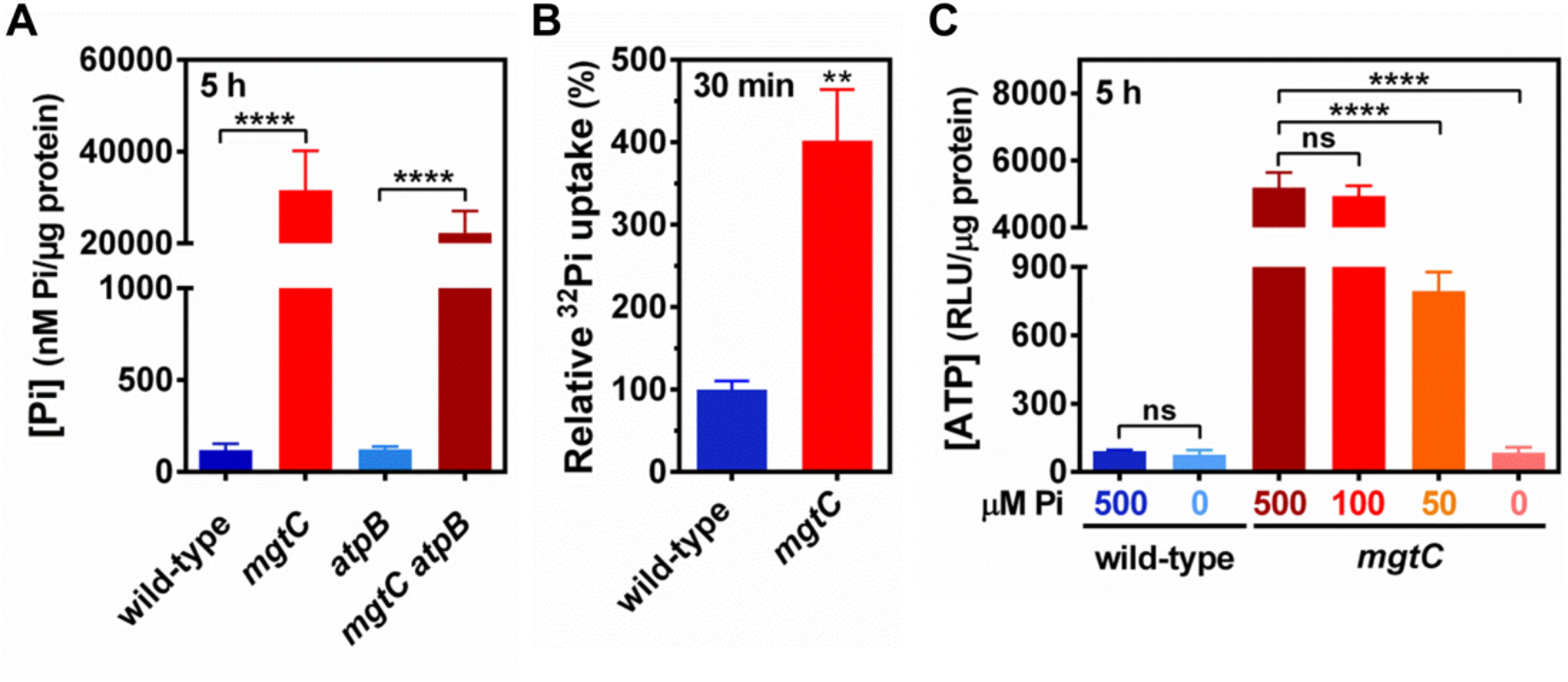
MgtC-dependent inhibition of Pi transport and assimilation into ATP during cytoplasmic Mg^2+^ starvation. *(A)* Total intracellular Pi in wild-type (14028s), *mgtC* (EL4), *atpB* (MP24) or *mgtC atpB* (MP25) *Salmonella* following 5 h of growth in MOPS medium containing 10 μM MgCl_2_ and 2 mM K_2_HPO_4_. *(B)* Relative radioactive orthophosphate (^32^Pi) uptake in wild-type (14028s) or *mgtC* (EL4) cells. Bacteria were grown in MOPS medium containing 10 μM MgCl_2_ and 500 μM K_2_HPO_4_ during 3 h before the addition of ^32^Pi to the cultures. Levels of ^32^Pi accumulated in cells were determined after 30 min of labeling by liquid scintillation counting, as described in Materials and Methods. ^32^Pi uptake values were normalized against the wild-type strain. *(C)* Intracellular ATP levels in wild-type (14028s) or *mgtC* (EL4) *Salmonella* following 5 h of growth in MOPS medium containing 10 μM MgCl_2_ and the indicated concentration of K_2_HPO_4_ (μM Pi). Means ± SDs of at least three independent experiments are shown. **P < 0.01, ****P < 0.0001, and ns, no significant difference. Two-tailed *t* tests *(A-C)*.

To test the aforementioned hypothesis, we measured transport of radiolabeled Pi (^32^Pi) following MgtC expression from its native chromosomal location in response to cytoplasmic Mg^2+^ starvation. We established that *mgtC* cells accumulated four times more radioactivity than the isogenic wild-type strain after 30 min of growth in the presence of ^32^Pi (Fig. 2*B*). This indicated that the increased steady-state intracellular Pi levels observed for *mgtC* and *mgtC atpB* mutants (Fig. 2*A*) arises from the uptake of extracellular Pi, as opposed to increased Pi release from intracellular sources. [Note that all ^32^Pi transport assays were performed in the presence of cold Pi, at a molar ratio of 1:25 (^32^Pi:Pi), to prevent the expression of the PstSCAB Pi transporter resulting from lack of Pi in the growth medium (1). The influx of Pi observed in the assays is underestimated (see Discussion and Materials and Methods)].

If MgtC functions to restrict cytoplasmic Pi availability, and Pi is required for ATP synthesis, we posited that the increased ATP levels in an *mgtC* mutant, but not in wild-type *Salmonella*, could be regulated by the availability of Pi in the growth medium. To test this prediction, we measured ATP levels in wild-type and *mgtC* cells grown in minimal media with low Mg^2+^ and decreasing concentrations of exogenous Pi. [Note that bacteria are able to grow in medium lacking exogenous Pi, due to the residual Pi content present in the mixture of casamino acids supplemented to the culture medium (see Materials and Methods)]. Strikingly, we established that, during cytoplasmic Mg^2+^ starvation, the ATP levels in an *mgtC* mutant could be reduced by decreasing exogenous Pi from 500 to 0 μM (Fig. 2*C*). By contrast, ATP levels in wild-type cells remained invariably low (Fig. 2*C*). Importantly, in the absence of exogenous Pi, wild-type and *mgtC* cells had similar levels of ATP (Fig. 2*C*). Taken together, these results indicate that MgtC controls ATP synthesis by limiting cellular Pi uptake.

### MgtC inhibits a non-canonical Pi transport system

*Salmonella enterica* encodes two *bona fide* Pi import systems: *pitA* and *pstSCAB* (Fig. 3*A*). While PitA functions as a metal:phosphate (M:Pi)/proton symporter (1, 49–51), PstSCAB works as a high affinity, ATP-dependent Pi transporter (Fig. 3*A*) (1, 6, 51, 52). In addition to *pitA* and *pstSCAB, Salmonella* also harbors an *yjbB* homolog, which encodes a sodium/phosphate symporter. Whereas YjbB has been shown to promote Pi export (53), we performed experiments under a cautious assumption that this protein may also be able to import Pi (Fig. 3*A*).

**Figure 3.**
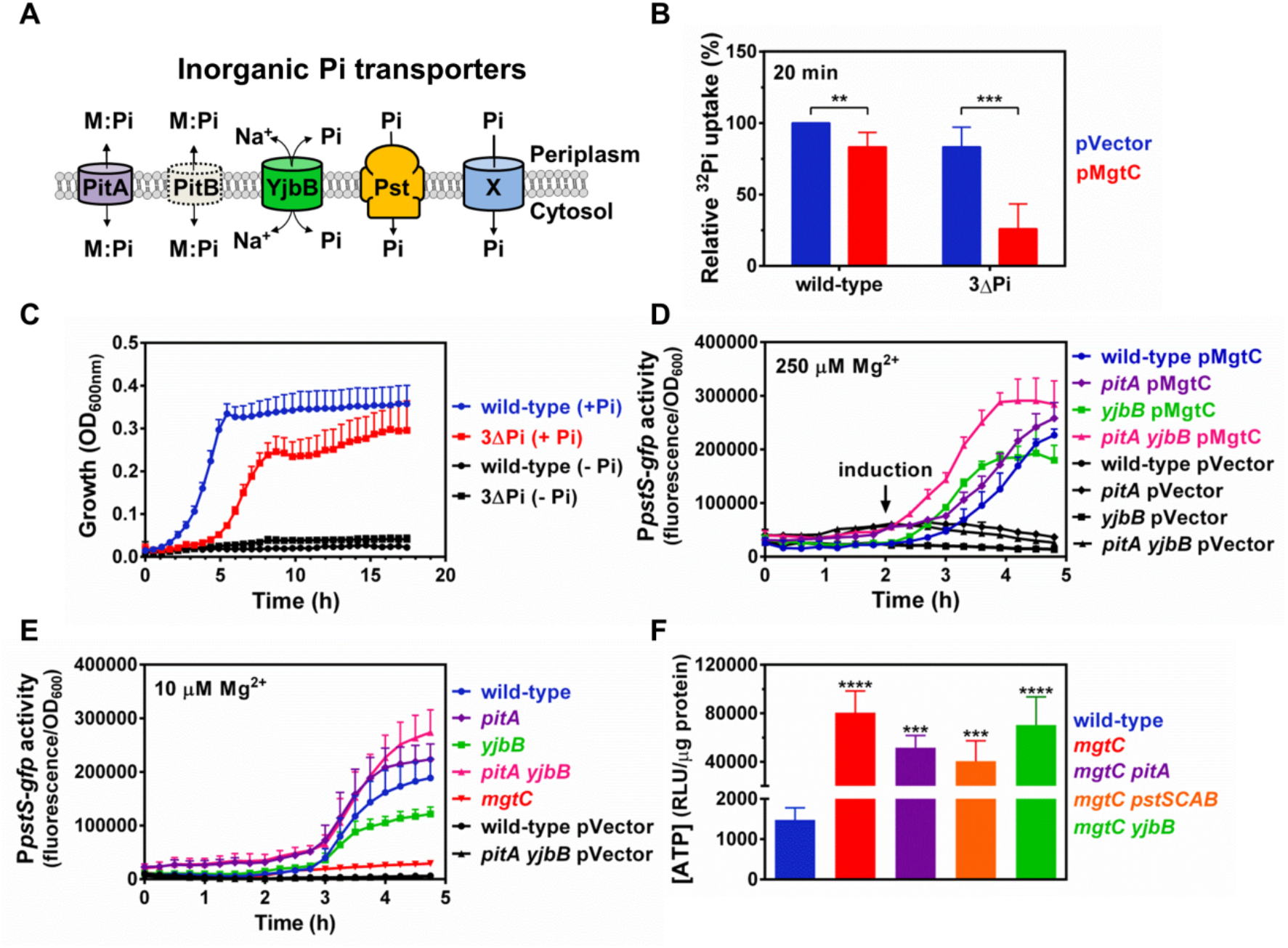
A non-canonical Pi transport system is inhibited by MgtC. *(A)* Schematic representation of inorganic Pi transporters harbored by *Salmonella enterica* and *Escherichia coli*. Note that PitB (grey, dashed outline) is absent from *Salmonella.* M:Pi, metal-phosphate complex; X, inferred, uncharacterized Pi transporter inhibited by MgtC. *(B)* Relative ^32^Pi uptake in wild-type (14028s) or Δ3Pi (RB39) carrying either pVector (pUHE-21) or pMgtC (pUHE-MgtC). Bacteria were grown in MOPS medium containing 250 μM MgCl_2_ and 500 μM K_2_HPO_4_ until OD_600_ ≈ 0.2. Cultures were then propagated for 15 min in the presence of 750 μM isopropyl β-d-1-thiogalactopyranoside (IPTG) prior to the addition of ^32^Pi. Transport of ^32^Pi was allowed to take place for 20 min. Intracellular ^32^Pi accumulation was determined by liquid scintillation counting, as described in Materials and Methods. ^32^Pi uptake values were normalized against the wild-type pVector strain. **P < 0.01, ***P < 0.001, unpaired two tailed *t* test. *(C)* Growth curve of wild-type (14028s) or Δ3Pi (RB39) *Salmonella.* Cells were grown in MOPS medium uyand containing 10 mM MgCl_2_ and either 0 (-Pi) or 500 (+Pi) μM of K_2_HPO_4_. *(D)* Fluorescence from wild-type (14028s), *pitA* (MP1251), *yjbB* (MP1252), *pitA yjbB* (MP1479p) *Salmonella* carrying *pPpstS-* GFPc and either pVector or pMgtC. Cells were grown in MOPS medium containing 250 μM MgCl_2_ and 500 μM K_2_HPO_4_. 250 μM of IPTG were added after 2 h of growth. *(E)* Fluorescence from wild-type (14028s), *pitA* (MP1251), *yjbB* (MP1252), *pitA yjbB* (MP1479p), *mgtC* (EL4) *Salmonella* carrying *pPpstS-* GFPc or pVector (the promoterless GFP plasmid pGFPc). Cells were grown in MOPS medium containing 10 μM MgCl_2_ and 500 μM K_2_HPO_4_. *(F)* Intracellular ATP levels in wild-type (14028s), *mgtC* (EL4), *mgtC pitA* (MP1254), *mgtC pstSCAB* (MP1720) or *mgtC yjbB* (MP1255) *Salmonella* following 5 h of growth. Cells were grown in MOPS medium containing 10 μM MgCl_2_ and 2 mM K_2_HPO_4_. ***P < 0.001, ****P < 0.0001, unpaired two-tailed *t* tests against the wild-type strain. For all graphs (*B-F*), means ± SDs of at least three independent experiments are shown.

To directly test the role of MgtC on the inhibition of PitA, PstSCAB or YjbB, we measured ^32^Pi uptake following ectopic MgtC expression in the wild-type and a *pitA pstSCAB yjbB* triple mutant (3ΔPi) strains. In wild-type cells, harboring all known Pi transport systems, MgtC expression led to a mild (17%) decrease in ^32^Pi uptake when compared with the empty vector control (Fig. 3*B*). By contrast, ectopic expression of MgtC reduced Pi uptake by 70% in the 3ΔPi background, relative to the vector control (Fig. 3*B*), indicating that MgtC decreases Pi influx in a *pitA pstSCAB yjbB-* independent manner. In support of these results, a 3ΔPi is able to grow using Pi as the sole P source (Fig. 3*C*), indicating that a yet unidentified transporter imports Pi into the cytoplasm to support the growth of this *Salmonella* strain.

Two additional lines of evidence indicated that MgtC inhibits the activity of a non-canonical Pi transporter. First, because MgtC expression causes a shortage in cytoplasmic Pi (Fig. 2 and 3*B*), ectopic expression of MgtC elicits a dose-dependent activation of the PhoB/PhoR two-component system and transcription of the PhoB-activated PstSCAB transporter (Fig. S1) (15). Hence, deletion of the MgtC-targeted Pi transporter should result in a constitutively high *pstS-gfp* activity that is irresponsive to MgtC expression. In other words, deletion of the Pi transporter should be epistatic to the effect of MgtC expression on PhoB/PhoR activation levels. Consistent with the notion that MgtC inhibits a non-canonical Pi transporter, we determined that expression of MgtC (either ectopically in medium containing 250 μM Mg^2+^ or natively in medium containing 10 μM Mg^2+^) induced *pstS* transcription in *pitA* and *yjbB* single and double mutant strains (Fig. 3*D-E*). [We purposely did not carry out this epistatic analysis with the *pstSCAB* mutant for two reasons. First, the PstSCAB transporter participates in the regulation of the PhoB/PhoR two-component system through a physical interaction (54). Consequently, inactivation of the transporter through mutations in *pst* genes leads to PhoB/PhoR hyperactivation, effectively preventing signal transduction (1, 3, 54, 55). Second, we reasoned that it would be unlikely for cells to have evolved a regulatory circuit whereby MgtC inhibition of PstSCAB transporter activity would promote PstSCAB expression (Fig. S1) (15), requiring more MgtC protein, *ad infinitum*]. Second, we posited that deleting the MgtC-inhibited transporter in an *mgtC* background should abolish its exacerbated ATP accumulation (Fig. 2*C*). However, mutations in either *pitA*, *pstSCAB* or *yjbB* did not lower the intracellular ATP levels of an *mgtC* mutant (Fig. 3*F*). In sum, these experiments indicate that MgtC inhibits Pi uptake by targeting an unidentified Pi transporter.

### Phosphate limitation rescues the translation and growth defects of an *mgtC* mutant experiencing cytoplasmic Mg^2+^ starvation

The increased ATP levels in an *mgtC* mutant experiencing cytoplasmic Mg^2+^ starvation causes ribosomal assembly defects, presumably because Mg^2+^ ions required for ribosome stabilization are bound to ATP molecules (26). This results in inefficient translation, growth arrest and a loss of viability (19, 26), all of which can be reversed by enzymatic hydrolysis of ATP (26, 27). Given that MgtC lowers ATP synthesis by inhibiting Pi acquisition (Fig. 2*A*, 2*C* and 3*B*), we reasoned that the translation, the growth defects and the loss of viability of an *mgtC* mutant would also be rescued by limiting the access of cells to Pi. To test this notion, we measured translation rates in the wild-type and *mgtC* strains experiencing cytoplasmic Mg^2+^ starvation in the presence of 500 or 0 μM exogenous Pi. As predicted, wild-type and *mgtC* strains showed similar translation rates in the absence of exogenous Pi (Fig. 4*A*-*B*). By contrast, during growth at 500 μM Pi, the translational rate of the *mgtC* mutant was 4.7-fold reduced in comparison with wild-type levels (Fig. 4*A*-*B*).

**Figure 4.**
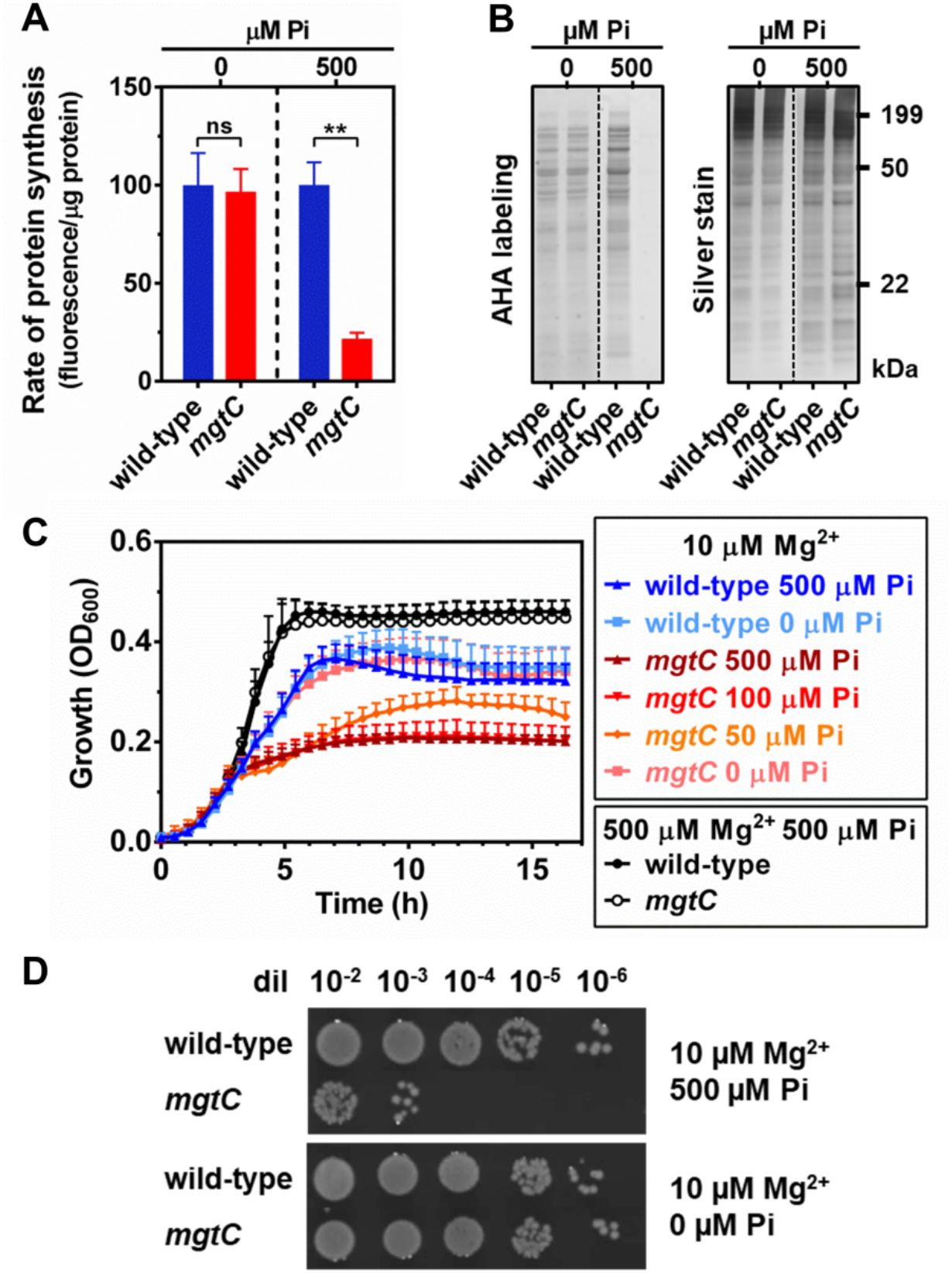
Effect of phosphate limitation on the translation rate, growth and viability of an *mgtC* mutant during cytosolic Mg^2+^ starvation. *(A)* Quantification and *(B)* SDS-PAGE analysis of the rate of protein synthesis [L-azidohomoalanine (AHA) labeling] of wild-type or *mgtC* (EL4) *Salmonella.* Cells were grown in MOPS medium containing 10 mM MgCl_2_ and 2 mM K_2_HPO_4_ until OD_600_ ≈ 0.4. Cells were subsequently washed thrice with MOPS medium lacking MgCl_2_, K_2_HPO_4_ and amino acids, and grown for additional 90 min in MOPS medium lacking methionine, and containing 10 μM MgCl_2_ plus the indicated concentration of K_2_HPO_4_ (μM Pi). Means ± SDs of four independent experiments are shown. The gel is representative of four independent experiments. Samples from 0 and 500 μM Pi were resolved and imaged from different gels (indicated by dashed lines). *(C)* Growth curve of wild-type (14028s) or *mgtC* (EL4) *Salmonella.* Cells were grown in MOPS medium containing the indicated concentrations of MgCl_2_ and K_2_HPO_4_. Means ± SDs of three independent experiments are shown. *(D)* Viable cell count of wild-type (14028s) or *mgtC* (EL4) *Salmonella* following 16 h of growth in MOPS medium containing 10 μM MgCl_2_ and 0 or 500 μM K_2_HPO_4_. Cell suspensions were normalized to the same OD_600_, diluted, and 5 μL were spotted on plates. Images were taken after incubation of plates at 37°C for 18 h, and are representative of three independent experiments.

Next, we measured the effect of reducing exogenous Pi on growth and viability of strains grown in low Mg^2+^ medium. We established that steadily decreasing exogenous Pi in the growth medium from 500 to 0 μM progressively increased the growth of the *mgtC* mutant strain (Fig. 4*C*). Remarkably, in the absence of exogenous Pi, the *mgtC* mutant displayed growth yield and kinetics indistinguishable from that of the wild-type strain (Fig. 4*C*). Furthermore, during growth in medium lacking Pi, the loss of viability observed in the *mgtC* mutant (19) was suppressed, and the mutant maintained the same number of colony forming units (CFU) per optical density unit (OD_600_) as the wild-type strain (Fig. 4*D*). As expected, the growth of wild-type and *mgtC* mutant cultures in medium containing 500 μM Pi were indistinguishable when Mg^2+^ was made abundant by raising its concentration from 10 to 500 μM (Fig. 4*C*). Altogether, these results establish that MgtC functions to prevent cytotoxic effects resulting from excessive Pi uptake.

### Pi cytotoxicity results from its exacerbated assimilation into ATP and subsequent chelation of Mg^2+^ during growth under conditions of high Mg^2+^

Excessive Pi uptake has been known to hinder bacterial growth in conditions where access to extracellular Mg^2+^ is not limiting and, consequently, cells are not anticipated to experience cytoplasmic Mg^2+^ starvation (3, 7). For instance, in *E. coli*, mutations that increase PstSCAB activity or expression cause heightened Pi uptake and growth inhibition (1–4). Given the aforementioned experimental results, we hypothesized that excess Pi would exert its cytotoxicity following its incorporation into ATP, and subsequent chelation of essential cations, particularly Mg^2+^.

According to this hypothesis, PstSCAB overexpression should promote phenotypes caused by cytoplasmic Mg^2+^ starvation—i.e. elevated ATP levels, inhibition of translation and growth, and induction of MgtC expression (19, 25, 26). Additionally, these phenotypes should be rescued by increasing the availability of free cytoplasmic Mg^2+^, through the provision of excess Mg^2+^ in the growth medium, or the reversal of P assimilation through enzymatic hydrolysis of ATP (19, 26, 27). To test this notion, we initially measured ATP levels and growth of wild-type cells following ectopic over-expression of the PstSCAB transporter from a plasmid. During growth in medium containing high (10 mM) Pi and intermediate (0.1 mM) Mg^2+^ levels, cells do not typically experience cytoplasmic Mg^2+^starvation (20, 21) and, consequently, do not express MgtC (Fig. 6*A*). Under these growth conditions, PstSCAB overexpression caused approximately a 4-fold increase in ATP levels in comparison to control strains harboring either an empty vector or a plasmid expressing the inner membrane protein PmrB (Fig. 5*A*). Notably, this rise in ATP levels was accompanied by reductions in growth rate and growth yield, two phenotypes not observed in the control strains (Fig. 5*B*).

**Figure 5.**
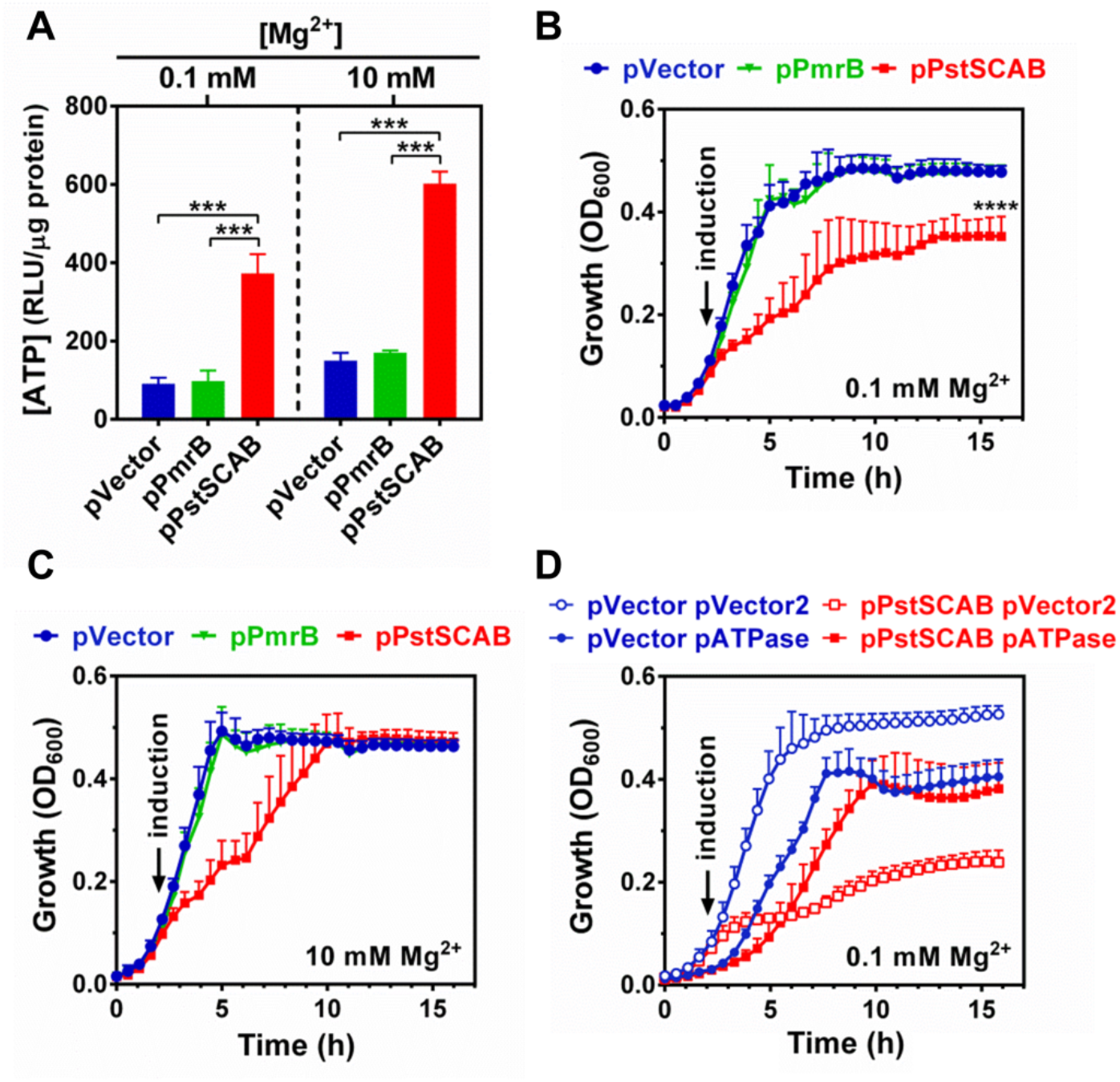
Effect of PstSCAB on growth, P assimilation and Mg^2+^ homeostasis. *(A)* Intracellular ATP levels of cultures depicted in *(B-C).* Measurements were conducted at 5 h of growth. *(B-C)* Growth curves of wild-type (14028s) *Salmonella* carrying pVector (pUHE-21), pPstSCAB (pUHE-PstSCAB), or pPmrB (pUHE-PmrB) in MOPS medium containing 0.1 (*B*) and 10 mM MgCl_2_ (*C*). *(D)* Growth curve of wild-type (14028s) *Salmonella* carrying either pVector or pPstSCAB, and either pVector2 (pBbB2K-GFP) or pATPase (pBbB2K-AtpAGD). In all experiments, cells were grown in MOPS medium containing 10 mM K_2_HPO_4_ and the indicated MgCl_2_ concentration. Ectopic protein expressions were induced by adding 250 μM IPTG to the cultures after 2 h of growth. Means ± SDs of three independent experiments are shown. *(A-B)* ***P < 0.001, ****P < 0.0001, unpaired two tailed *t* test against pVector.

**Figure 6.**
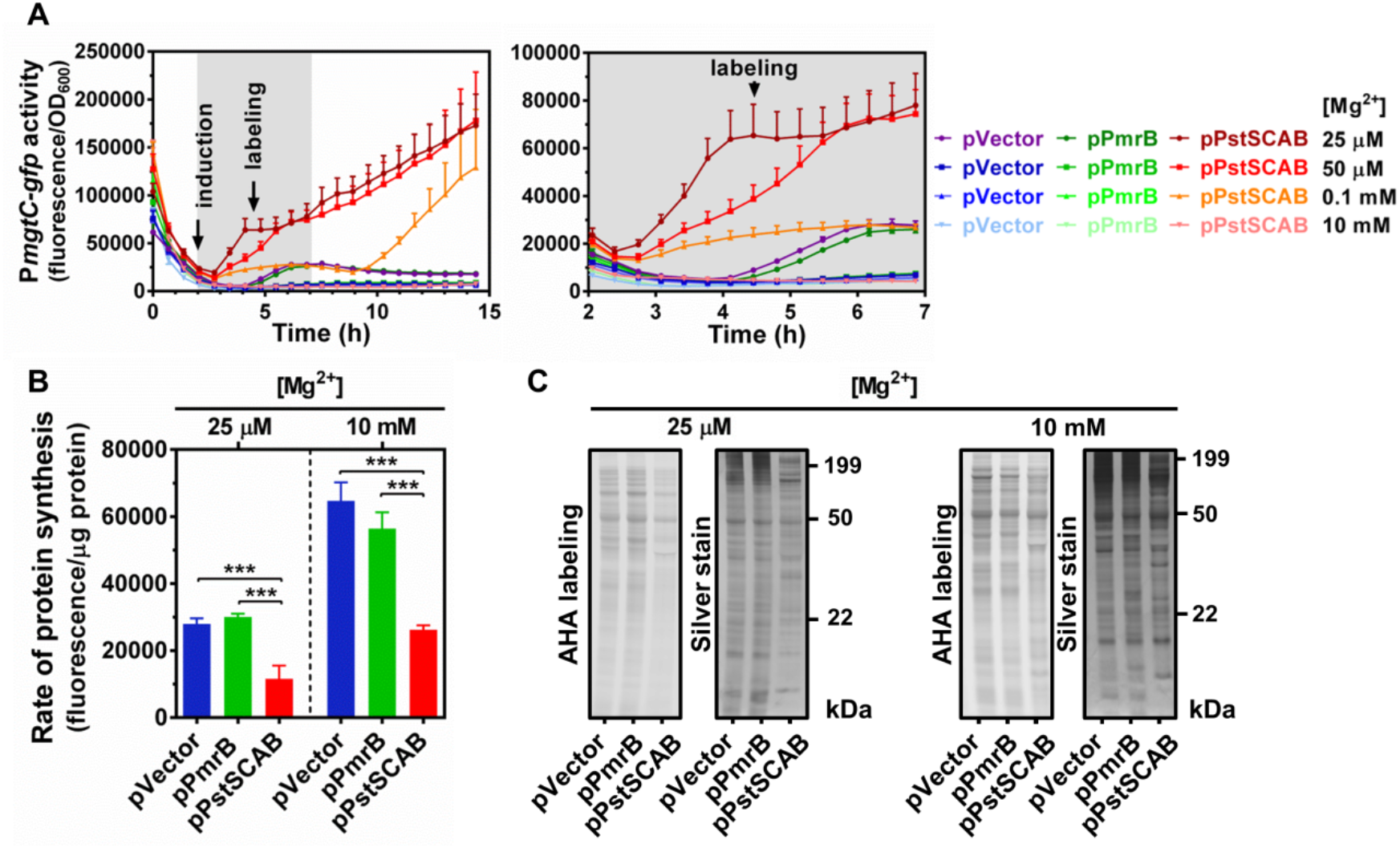
Effect of PstSCAB expression on *mgtC* transcription and cellular translation rates during grown under conditions of moderate and high Mg^2+^. *(A)* Fluorescence from wild-type (14028s) *Salmonella* carrying transcriptional reporter pP*mgtC*-GFPc and either pVector (pUHE-21), pPstSCAB (pUHE-PstSCAB), or pPmrB (pUHE-PmrB). Full time course (left hand-side panel) and inset between 2 and 7 h (right hand-side panel) of the experiments are shown. *(B)* Quantification and *(C)* SDS-PAGE analysis of the rate of protein synthesis [L-azidohomoalanine (AHA) labeling] of wild-type (14028s) *Salmonella* carrying either pVector, pPmrB or pPstSCAB. In all experiments, cells were grown in MOPS medium containing 10 mM K_2_HPO_4_ and the indicated concentration of MgCl_2_. For AHA labeling, bacteria were cultured in MOPS medium lacking methionine (see Materials and Methods). 250 μM IPTG was added to the cultures following 2 h of growth. AHA was incorporated to the cultures at 4.5 h. Means ± SDs of three independent experiments are shown. Gels are representative of three independent experiments. ***P < 0.001, unpaired two tailed *t* test against pVector.

Two lines of evidence indicated that the growth defect resulting from PstSCAB-induction was caused by an increase in ATP and chelation of free Mg^2+^. First, in the presence of a 100-fold excess of Mg^2+^ (10 mM), PstSCAB expression caused a minor reduction in growth rate, but enabled cells to reach the same growth yields as the control strains (Fig. 5*C*). Importantly, Mg^2+^ rescued the growth of the PstSCAB expressing strain even though its ATP levels remained 4-fold higher relative to control strains harboring the empty vector or expressing the plasmid-borne PmrB (Fig. 5*A*). Second, ectopic expression of a plasmid-encoded ATPase (15) rescued the reduction in growth yield caused by PstSCAB-induction (Fig. 5*D*). By contrast, no rescue was observed in cells harboring the vector control (Fig. 5*D*). Taken together, these results indicate that excess Pi imported into the cytoplasm is rapidly assimilated into ATP, causing a decrease in the levels of free cytoplasmic Mg^2+^, and inhibiting growth.

### Excessive Pi uptake impairs translation and promotes MgtC expression during growth in high Mg^2+^

If excessive Pi uptake leads to physiological conditions resembling cytoplasmic Mg^2+^ starvation (Fig. *5A-D*) (19, 26, 27), then, increased Pi uptake resulting from the expression of PstSCAB should also decrease translation rates. Furthermore, because a reduction in translation rates caused by cytoplasmic Mg^2+^ starvation promotes *mgtC* transcription (20, 21, 25, 26, 38), excessive Pi uptake should promote MgtC expression.

To test these predictions, we first measured fluorescence over time in otherwise wild-type strains harboring a plasmid-borne *gfp* transcriptional fusion to the promoter and leader regions of *mgtC*, and either an empty vector, the pPmrB, or the pPstSCAB plasmid. We determined that *mgtC-gfp* fluorescence increased one hour following the induction of PstSCAB expression (Fig. 6*A*). This increase in fluorescence was absent or delayed in control strains harboring the empty vector or expressing the plasmid copy of the inner membrane protein PmrB (Fig. 6*A*). [Note that during growth in 25 μM Mg^2+^, the control strains display a minor increase in fluorescence at 4 h of growth. This small increase in fluorescence results from late onset of cytoplasmic Mg^2+^, which is delayed when compared to cells grown at 10 μM Mg^2+^ (15, 20, 21, 56)]. Consistent with the notion that increased ATP production resulting from excessive Pi assimilation disrupts translation by sequestering free Mg^2+^ ions, the effect of PstSCAB expression on the time and fluorescence levels of *mgtC-gfp* was inversely related to the availability of Mg^2+^ in the growth medium. Specifically, fluorescence levels resulting from PstSCAB expression in cultures grown in 25 μM Mg^2+^ were higher than those grown in 50 μM Mg^2+^, which, in turn, were higher than those grown in 100 μM Mg^2+^ (Fig. 6*A*). Whereas cells grown in 100 μM Mg^2+^ displayed a relatively mild increase in *mgtC-gfp* activity 1 h after PstSCAB expression, their fluorescence levels rose rapidly at 7.5 h post induction (Fig. 6*A*). This later induction of *mgtC* expression likely reflects the time point at which cells exhaust the Mg^2+^ available in the growth medium, can no longer neutralize the excess of intracellular Pi incorporated into ATP and, consequently, are unable to efficiently stabilize their ribosomes (26).

To directly test if excessive Pi uptake affected ribosome activity, we measured the translation rates of the aforementioned strains following the induction of the plasmid-borne proteins. We determined that during growth in medium containing 25 μM Mg^2+^, PstSCAB expression caused a 2-fold reduction in translation rates relative to control strains (Fig. 6*B-C*). Notably, PstSCAB expression also caused a relative reduction in translation rates during growth in medium containing a 400-fold excess (10 mM) of Mg^2+^ (Fig. 6*B-C*). However, at 10 mM Mg^2+^, cellular translation rates were approximately 2-fold higher across all strains (Fig. 6*B-C*). Taken together, these results indicate that the rapid assimilation of excessive Pi imported in the cytoplasm reduces levels of free cytoplasmic Mg^2+^, thereby lowering translation efficiency and promoting MgtC expression.

## Discussion

### Pi toxicity via disruption of Mg^2+^ homeostasis

During cytoplasmic Mg^2+^ starvation, *Salmonella* expresses the MgtC membrane protein. This protein promotes bacterial growth and viability by virtue of its ability to reduce ATP levels, thereby increasing the concentration of free intracellular Mg^2+^ needed for the functioning of vital Mg^2+^-dependent processes (19, 21, 25–27). In this work, we demonstrate that MgtC lowers ATP levels by inhibiting Pi uptake (Fig. 2*A-C* and 3*B*), thus limiting the availability of an ATP precursor rather than interfering with the enzymatic catalysis of ATP-generating reactions *per se* (Fig. 1*A-C* and 2*A-C*). We establish that MgtC hinders the activity of a yet unidentified transporter, which functions as the main Pi uptake system in *Salmonella* (Fig. 3*A-F*). Finally, we show that, even when cytoplasmic Mg^2+^ is not limiting, excessive Pi uptake leads to increased ATP synthesis, depletion of free cytoplasmic Mg^2+^, inhibition of translation and growth (Fig. 5*A-D* and 6*A-C*). These results indicate that bacterial cells control Pi uptake and subsequent assimilation to avoid the depletion of free cytoplasmic Mg^2+^.

The capacity of MgtC to maintain physiological ATP levels during cytoplasmic Mg^2+^ starvation had been so far ascribed to its inhibitory effect on *Salmonella’s* F_1_F_o_ ATP synthase (30). Here, we demonstrate that this inaccurate conclusion resulted from an experimental artifact arising from the propagation of *atpB* strains in a poorly fermentable carbon source (Fig. 1*A-C*). Indeed, the notion that MgtC lowers ATP levels in an F_1_F_O_ ATP synthase-independent fashion has found support on a recent study by an independent group. While also growing cells in a poorly fermentable carbon source, the authors were still able to show that overexpression of MgtC lowers ATP levels in *atpB* mutants (40). Yet, while this study was unable to identify the source of ATP reduction, we now determine its origin. We establish that MgtC limits Pi uptake, simultaneously hindering *all* ATP-generating enzymatic reactions in the cell (Fig. 7).

**Figure 7.**
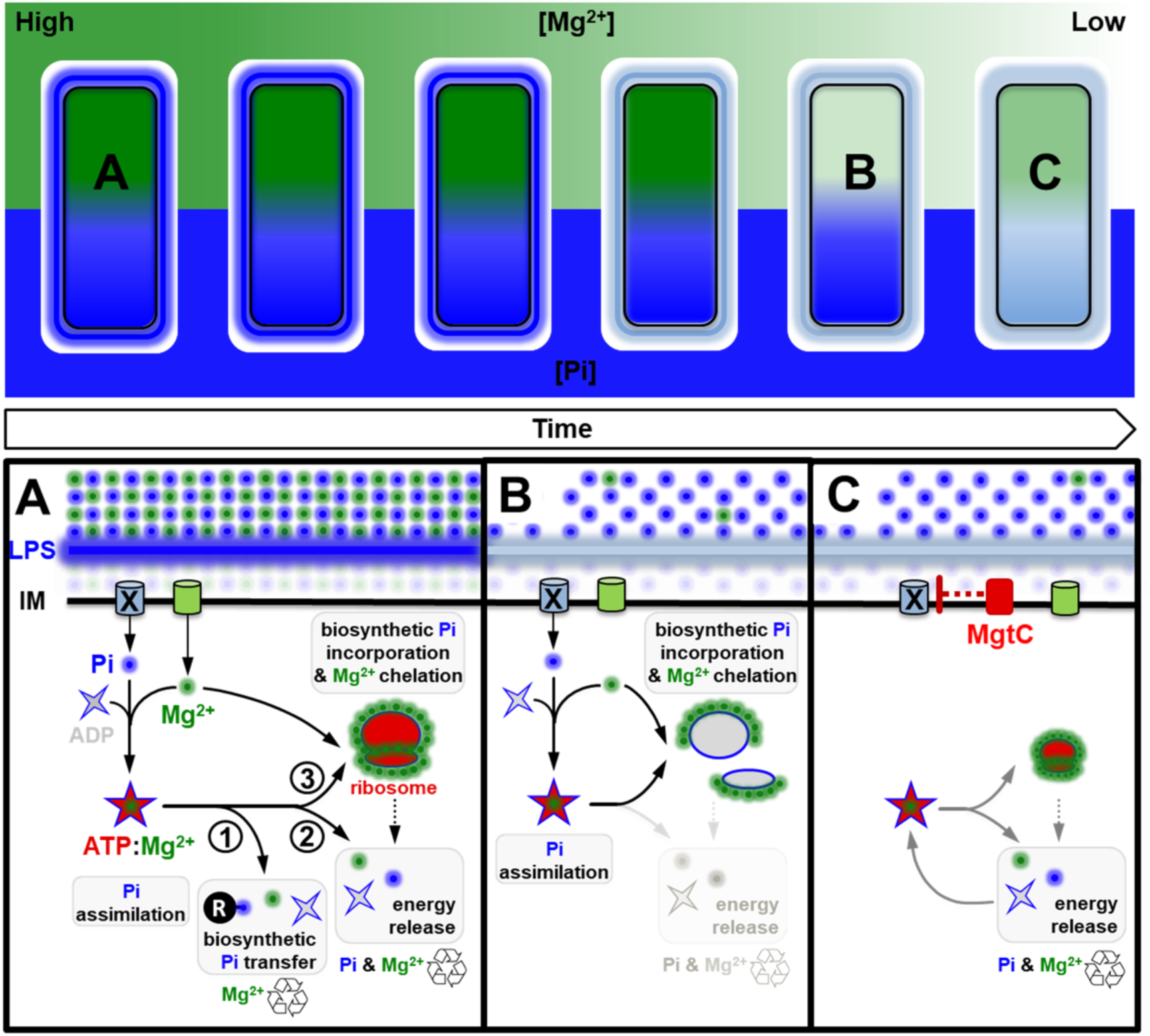
Model illustrating how limitation of phosphate assimilation maintains cytoplasmic Mg^2+^ homeostasis in *Salmonella enterica.* Top panel: Overview of the temporal adaptation of *Salmonella enterica* to Mg^2+^ starvation. Mg^2+^ (green) and Pi (blue) concentrations in the environment, lipopolysaccharide (LPS) and cytoplasm are depicted as gradients with dark colors denoting high concentration and light colors representing low concentrations. Bottom panel: Schematic depicting molecular events and responses underlying the adaptation. *(A)* During homeostasis, Pi is imported into the cytoplasm through dedicated inner membrane (IM) transport systems [X (unknown transporter) and PitA]. Cells assimilate imported Pi through the synthesis of ATP, which exists as a salt with positively charged Mg^2+^ (ATP:Mg^2+^). ATP:Mg^2+^ mediates the (1) transfer of Pi among biological molecules, (2) powers energy-dependent enzymatic reactions, and (3) promotes ribosome biogenesis (1). ATP hydrolysis for the release of energy recycles Pi and Mg^2+^, whereas the biosynthetic transfer of Pi typically recycles cytoplasmic Mg^2+^. *(B)* After consuming the Mg^2+^ present in the environment, cells eventually experience a shortage in cytoplasmic Mg^2+^ levels (20, 21, 56). Insufficient cytoplasmic Mg^2+^ impairs ribosomal subunit assembly, lowering translation efficiency (26). This reduces the consumption of ATP by translation reactions and, consequently, decreases the recycling of Mg^2+^ and Pi from ATP:Mg^2+^ (15). (*C*) *Salmonella enterica* expresses the MgtC membrane protein as a homeostatic response to cytoplasmic Mg^2+^ starvation. MgtC inhibits Pi uptake through an unknown transporter (X), thereby preventing assimilation Pi into ATP. As the levels of ATP and ribosomes decrease, free Mg^2+^ ions necessary for core processes such as translation are recovered, increasing the efficiency of protein synthesis, and the recycling of Mg^2+^ and Pi from ATP:Mg^2+^. The density and size of cartoons represent the concentrations of Mg^2+^, Pi, and ribosomes.

MgtC inhibition of Pi uptake occurs independently of all characterized Pi importers (PitA and PstSCAB) as well as a Pi exporter (YjbB). Interestingly, heterologous MgtC expression in *E. coli* also activates its PhoB/PhoR two-component system (Fig. S2). Three bona fide Pi importers have been described in *E. coli:* PstSCAB, PitA and PitB, which is not present in *Salmonella* (Fig. 3*A*) (1, 51, 57). Notably, *E. coli* strains containing mutations in all these three Pi transport systems are still able to grow on minimal medium with Pi as the only P source (51, 58). This indicates that *E. coli* also encodes a homolog of the Pi transporter that is targeted by MgtC in *Salmonella*. In fact, given that MgtC homologs promote growth in low Mg^2+^ in a large number of distantly related species (32, 59–65) this transporter is likely to be wide spread in bacteria.

If MgtC inhibits the activity of a single Pi importer, what then, prevents Pi uptake by other transport systems? Transcription of the high affinity PstSCAB Pi importer is induced by the PhoB/PhoR two-component system in response to a decrease in cytoplasmic Pi. During the initial stages of cytoplasmic Mg^2+^ starvation, the ribosomal subunits are unable to assemble efficiently (26). This leads to a decrease in translation efficiency and a concomitant reduction in ATP hydrolysis and free cytoplasmic Pi, which triggers *pstSCAB* transcription (15). Whereas expression of MgtC leads to inhibition of the main Pi transporter, the expression of MgtA and MgtB results in an influx of Mg^2+^ into the cytoplasm (19–24). This response restores ribosomal subunit assembly and increases translation efficiency (Fig. 7). Efficient ATP consumption by translation reactions (most notably the charging of tRNAs and the synthesis of GTP, which is subsequently used by elongation factors) (26), replenishes intracellular Pi, effectively repressing PhoB/PhoR and *pstSCAB* transcription (15).

Transcription of *pitA* is regulated by the availability of Pi:Zn^2+^ salt (66). Yet, in *E. coli,* PitA undergoes post-transcriptional repression during Mg^2+^ starvation. This repression is orchestrated by the Mg^2+^-sensing PhoP/PhoQ two-component system (67), which, in *Salmonella*, activates transcription of *mgtCB* and *mgtA* (19). Interestingly, growth in low Mg^2+^ decreases *pitA* mRNA levels in both *Salmonella* and *Yersinia pestis* (68, 69). This suggests that inhibition of PitA expression is a common feature of the Mg^2+^ starvation response among enteric bacteria. Because PitA transports M:Pi salts, repression of this gene may also prevent the importation of other metal cations, such as zinc, that can readily replace scarce Mg^2+^ in enzymatic reactions (70–72). In this context, it is interesting to note that the Pho84 Pi transporter of *Saccharomyces cerevisiae* can promote metal toxicity by importing M:Pi salts (73–76).

If excessive ATP is toxic during cytoplasmic Mg^2+^ starvation, why then does MgtC function by hindering Pi uptake and not by directly inhibiting ATP-generating enzyme(s)? Depending on the growth condition, the production of ATP via the oxidation of carbon can occur via several distinct pathways, each involving dozens of enzymes (48). While we can conceive that a single protein may have the capacity to directly inhibit a myriad of distinct enzymes, evolution has likely provided cells with a more parsimonious solution for this problem. Because Pi is an ATP precursor, limitation of Pi uptake enables cells to indirectly hinder *all* ATP generating reactions, *independently* from the metabolic pathway(s) used to oxidize the available carbon source(s) (Fig. 1*B* and 7). Hence, the inhibition of Pi uptake by MgtC allows *Salmonella* to lower ATP synthesis, eliciting a physiological response to a lethal depletion of cytoplasmic Mg^2+^, whether carbon is being metabolized via classic glycolysis, the Entner-Doudoroff, the Pentose Phosphate, the Tricarboxylic Acid Cycle or any other energy-generating pathway (Fig. 1*B* and 7).

### On the activation of PhoR by MgtC

The results presented herein challenge a proposed model of MgtC functionality set forth by Choi *et al*. (77). In this study, the authors hypothesize that MgtC promotes PstSCAB expression and Pi-uptake via the activation of the PhoB/PhoR two-component system through a direct physical interaction between MgtC and the PhoR histidine kinase (77). Several pieces of our data dispute this model and demonstrate that MgtC inhibits Pi uptake to maintain cellular viability during cytoplasmic Mg^2+^ starvation.

First, the growth and viability defects of an *mgtC* mutant are due to excess Pi uptake, which is acutely toxic to cells experiencing cytoplasmic Mg^2+^ starvation (Fig. 4*C-D*; see discussion above). Hence, it is hard to envision a model (77) where MgtC would promote the uptake of precisely the compound responsible for growth inhibition and loss of viability in cells lacking *mgtC* (Fig. 4*C-D*). In this light, there are discrepancies that arise in the model proposed by Choi *et al*. due to the lack of both physiological explanations and phenotypic assays to support the conclusion that cells import Pi during cytoplasmic Mg^2+^ starvation (77).

Second, when MgtC is expressed from its normal chromosomal location in response to cytoplasmic Mg^2+^ starvation, PhoB/PhoR activation and *pstSCAB* transcription precede MgtC expression, which persists long after *pstSCAB* transcription is subdued (Fig. S3*A-B*) (15). Hence, during physiologically relevant conditions, MgtC is unlikely to promote *pstSCAB* expression and Pi uptake as proposed (77). Instead, this phenomenon is better explained by the translation defect caused by a transient decrease in free cytoplasmic Mg^2+^, which is normalized by the activities of MgtA, MgtB and MgtC, expressed after PhoB/PhoR is activated (15, 26). Consistent with this notion, artificial influx of extracellular Pi by ectopic PstSCAB leads to increased ATP synthesis, disruption of free Mg^2+^ pools, inhibition of translation and promotion of *mgtC* transcription (Fig. 5*A-D* and 6*A-C*). This indicates that MgtC is expressed in response to stress(es) generated by increased Pi uptake. Notably, in *E. coli*, cytoplasmic Mg^2+^ starvation also disrupts translation homeostasis (26) and promotes a PhoB/PhoR activation (Fig. S4), even though this organism lacks an MgtC homolog.

Third, when Mg^2+^ is abundant, ectopic MgtC promotes *pstSCAB* expression (Fig. S1) (15). However, in contrast to results presented by Choi *et al.* (77) we established that increased *pstSCAB* transcription observed in this context *does not* lead to significant alterations in intracellular Pi levels (Fig. 3*B*). This is because, under these conditions, *pstSCAB* transcription is a homeostatic response resulting from the inhibition of a main Pi transporter by MgtC (Fig. 2*A-B* and 3*B-F*). These types of regulatory responses are common among multiple transport systems, and can be abolished by eliminating functional redundancy. For instance, while deletion of the house keeping Mg^2+^ transporter *corA* increases the expression of the specialized Mg^2+^ transporter MgtA (78), CorA overexpression repress MgtA production (56). That is, these genetic modifications alter the concentration of cytoplasmic Mg^2+^, which governs MgtA expression (20, 21). Likewise, deletion of the *pitA* Pi transporter (Fig. S5) or ectopic overexpression of MgtC leads to increased *pstS* transcription (Fig. S1). That is, cells sense and compensate for a shortage in intracellular Pi caused by these genetic alterations and increase expression of the PstSCAB system. In this light, ectopic MgtC expression in wild-type cells leads to a minor reduction in Pi uptake, but severely impairs Pi uptake in strains lacking the Pi transport systems encoded by *pitA*, *pstSCAB* and *yjbB* (Fig. 3*B*).

Fourth, substitution of PhoR leucine 421 by an alanine was reported to disrupt PhoR interaction with MgtC, preventing the activation of PhoB/PhoR by cytoplasmic Mg^2+^ starvation (77). We measured PhoB activation in wild-type and *phoR*^L421A^ strains using a *pstS-gfp* reporter fusion. Interestingly, wild-type and *phoR*^L421A^ strains displayed similar fluorescence levels, either when MgtC was ectopically induced (Fig. S6*A*), or when it was expressed from its normal chromosomal location during cytoplasmic Mg^2+^ starvation (Fig. S6*B*). Hence, PhoR L421 residue does not participate in a hypothetical MgtC-mediated PhoB/PhoR activation as proposed (77).

### Pi toxicity and the control of P assimilation

Mutations that increase Pi uptake via the PstSCAB transport system have been shown to inhibit growth in a wide range of bacterial species (1–7). Yet, the underlying molecular basis for this phenomenon has remained elusive. The realization that MgtC promotes growth and viability during cytoplasmic Mg^2+^ starvation (19, 21) by inhibiting Pi uptake (Fig. 2*A-B* and 3*B*) and, consequently, ATP synthesis (Fig. 2*C*) (27) prompted us to examine the physiological basis of Pi toxicity observed in the context of the aforementioned mutations. We established that increased Pi transport via PstSCAB causes a rise in ATP concentrations (Fig. 5*A*). Increased ATP disrupts the pools of free cytoplasmic Mg^2+^ (Fig. 6*A*), thereby inhibiting translation (Fig. 6*B-C*), growth (Fig. 5*B*), and precipitating the expression of MgtC when concentrations of Mg^2+^ in the growth medium is sufficiently high to silence its expression (Fig. 6*A*) (20, 21, 25). Whereas these results establish that the toxic effects of excessive Pi are manifested following its assimilation into ATP, they shed light into the underlying causes of Pi toxicity and reveal a logic for cellular control of P assimilation. Rapid synthesis of ATP, and ATP-derived highly charged Pi anions such as rRNA and poly-Pi (26, 79, 80), depletes the pools of free cytoplasmic Mg^2+^. Cells must control Pi uptake because, when biosynthetic precursors are abundant, simultaneous inhibition of all ATP-generating reactions in the cytoplasm cannot be easily attained.

## Materials and Methods

### Bacterial strains, plasmid constructs, primers, and growth conditions

The bacterial strains and plasmids used in this study are listed in Table S1, and oligonucleotide sequences are presented in Table S2. Single gene knockouts and deletions were carried out as described (81). Mutations generated via this method were subsequently moved into clean genetic backgrounds via phage P22-mediated transduction as described (82). For chromosomal point mutations, detailed strain construction is described below. Bacterial strains used in recombination and transduction experiments were grown in LB medium at 30° C or 37° C (81, 82). When required, the LB medium was supplemented with ampicillin (100 μg/mL), chloramphenicol (20 μg/mL), kanamycin (50 μg/mL), and/or L-arabinose (0.2% wt/vol).

Unless stated otherwise, physiological experiments with bacteria were carried out at 37° C with shaking at 250 rpm in MOPS medium (83) lacking CaCl_2_ (to avoid repression of the PhoP/PhoQ system) (84) and supplemented with 0.1% (w/v) bacto casamino acids (BD Difco), 25 mM glucose, and the indicated amounts of MgCl_2_ and K_2_HPO_4_. Experiments were conducted as follows: after overnight (~16-to 20-h) growth in MOPS medium containing 10 mM MgCl_2_ and 2 mM K_2_HPO_4_, cells washed three times in medium lacking Mg^2+^ and Pi and inoculated (1:100) in fresh medium containing the indicated concentrations of MgCl_2_ and K_2_HPO_4_ and propagated for the corresponding amount of time. It should be noted that at a concentration of 0.1% (w/v) bacto casamino acids (BD Difco), the medium already contains ~163 μM Pi. During physiological experiments, selection of plasmids was accomplished by the addition of ampicillin at 100 μg/mL (overnight growth) or 30 μg/mL (experimental condition), chloramphenicol at 20 μg/mL (overnight growth) or 10 μg/mL (experimental condition), and/or kanamycin at 50 μg/mL (overnight growth) or 20 μg/mL (experimental condition). Unless specified otherwise, heterologous expression of proteins was achieved by treatment of cultures with 250 μM (pMgtC, pPstSCAB) isopropyl β-D-1-thiogalactopyranoside (IPTG). ATPase expression from pBbB2k-AtpAGD was attained without the addition of the inductor.

### Estimation of intracellular ATP

Intracellular ATP was estimated as described (15). Briefly, luminescence measurements were performed in a SpectraMax i3x plate reader (Molecular Devices) with a BacTiter-Glo Microbial Cell Viability Assay Kit (Promega) in heat-inactivated cells (80° C for 10 min) according to the manufacturers instructions. Protein concentrations in cell samples were estimated using a Rapid Gold BCA Protein Assay Kit (Pierce). ATP measurements were normalized by the protein content of the samples.

### Estimation of intracellular Pi

Total Pi in the samples was estimated from crude cell extracts using the molybdenum blue method as described before (15, 85). The amounts of Pi in the samples were estimated from a standard curve generated from dilutions of a K_2_HPO_4_ solution of known concentration, and then normalized by the amount of protein present in each reaction.

### Phosphate transport assay

Wild-type (14028s) or *mgtC* (EL4) *Salmonella* were grown in MOPS medium containing 10 μM MgCl_2_ and 500 μM K_2_HPO_4_ during 3 h. Wild-type (14028s) or Δ3Pi (RB39) cells harboring either pVector or pMgtC were grown in MOPS containing 250 μM MgCl_2_ and 500 μM K_2_HPO_4_ until OD_600_ ≈ 0.2, at which point, MgtC expression was induced for 15 min with the addition of 750 μM IPTG. To assay the transport of Pi, 20 μCi of radioactive Pi solution (10 μL from a 2 mCi K_2_H^32^PO_4_ at a concentration of 2 mM of K_2_H^32^PO_4_, PerkinElmer cat. no. NEX055) was added to 1 mL of cell suspension. At the indicated time points, 50 μL of each sample was submitted to rapid filtration through 0.45 μm mixed cellulose ester membrane filters (Whatman) with an applied vacuum. The filters were washed three times with 1 mL of PBS buffer, and subsequently soaked in 5 mL of scintillation fluid (Research Products International). The amount of radioactivity taken up by the cells was determined with a scintillation counter (Triathler multilabel tester, HIDEX) using the ^32^P-window and by counting each vial for 20 s. Radioactive counts per minute were normalized by protein content using a Rapid Gold BCA Protein Assay Kit (Pierce). ^32^Pi uptake of each sample was normalized against the corresponding control in each independent experiment.

### Monitoring gene expression via fluorescence

Following overnight growth, bacteria were washed thrice and diluted in 1:100 in 1 mL of MOPS medium containing the appropriate concentrations of Mg^2+^ and Pi, and aliquot as technical replicates or triplicates into black, clear-bottom, 96-well plates (Corning). Two drops of mineral oil were used to seal the wells and prevent evaporation, and cultures were grown at 37° C with auto-mixing in a SpectraMax i3x plate reader (Molecular Devices). At the desired time points, the green fluorescence (excitation 485 nm/emission 535 nm) and absorbance at 600 nm (OD_600_) from the wells of the plates were read. Fluorescence measurements were normalized by the OD_600_ of the samples.

### Construction of plasmid pPstSCAB (pUHE-PstSCAB)

Phusion® High-Fidelity DNA Polymerase (New England Biolabs) was used in a PCR with primers 446 and 447 and *Salmonella* genomic DNA as template. The PCR product was resolved by agarose gel electrophoresis, purified using Monarch® Gel Extraction Kit (New England Biolabs), and ligated into BamHI/HindIII-digested pUHE-21-2-lacIq plasmid (86), using NEBuilder® HiFi DNA Assembly Cloning Kit (New England BioLabs). The assembly reaction was transformed into electrocompetent EC100D *E. coli*. The construct was verified by DNA sequencing using primers 448-452.

### Construction of plasmid pATPase (pBbB2K-AtpAGD)

Phusion® High-Fidelity DNA Polymerase (New England BioLabs) was used in a PCR with primers 799 and 800 and pUHE-AtpAGD (15) as template. PCR product was resolved by agarose gel electrophoresis, purified using Monarch® Gel Extraction Kit (New England Biolabs), and ligated into BamHI/EcoRI-digested pBbB2k-GFP plasmid (87), using NEBuilder® HiFi DNA Assembly Cloning Kit (New England BioLabs). The assembly reaction was transformed into electrocompetent EC100D *E. coli*. The functionality of the construct was tested by its capacity to decrease ATP levels in a *Salmonella mgtC* mutant strain (EL4) following growth in 10 μM Mg^2+^ medium (15).

### Construction of plasmid *pPmgtCB-GFP*

Phusion® High-Fidelity DNA Polymerase (New England Biolabs) was used in a PCR with primers W3443 and W3444 and plasmid pGFP303 (25) as template. The PCR product was resolved by agarose gel electrophoresis, purified using Monarch® Gel Extraction Kit (New England Biolabs), and ligated into SalI/HindIII-digested pACYC184 plasmid (88), using NEBuilder® HiFi DNA Assembly Cloning Kit (New England BioLabs). Assembly reactions were transformed into electrocompetent EC100D *E. coli*. The integrity of the construct was verified by DNA sequencing, and its functionality was verified by monitoring fluorescence in wild-type (14028s) *Salmonella* during growth in MOPS medium containing different MgCl_2_ concentrations (21).

### Construction of *phoR*^L421A^ strain (MP1665)

Phusion® High-Fidelity DNA Polymerase (New England Biolabs) was used in a PCR with primers 477 and 478 and plasmid pSLC-242 (89) as template. The PCR product was resolved by agarose gel electrophoresis, purified using Monarch® Gel Extraction Kit (New England Biolabs), and integrated into the chromosome of wild-type (14028s) *Salmonella* via λ-Red-mediated recombination using plasmid pSIM6 as described (81). Recombinant cells containing the insertion were selected on LB supplemented with 20 μg/mL chloramphenicol and 30 mM glucose at 30°C. This insertion was subsequently replaced via a second λ-Red mediated recombination of primer 480 into the chromosome. Cells were recovered for 3 h as described (89) and selected on MOPS medium containing 9.5 mM NH_4_Cl as the sole nitrogen source and 30 mM rhamnose as the sole carbon source. The identity of the *phoR*^L421A^ construct was verified by PCR with primers 481 and 482 followed by DNA sequencing. Its functionality was verified by introducing pP*pstS*-GFPc plasmid into this strain, and measuring fluorescence during growth in MOPS containing different K_2_HPO_4_ concentrations (15).

### L-azidohomoalanine (AHA) labeling and quantification

*Salmonella* strains were grown in MOPS medium supplemented with an amino acids mixture lacking methionine (Mix–Met: 1.6 mM of alanine, glycine, leucine, glutamate and serine, 1.2 mM glutamine and isoleucine, 0.8 mM arginine, asparagine, aspartate, lysine, phenylalanine, proline, threonine and valine, 0.4 mM histidine and tyrosine, and 0.2 mM cysteine and tryptophan), with the indicated MgCl_2_ and K_2_HPO_4_ concentrations. At the corresponding timepoints, cultures were labeled with 40 μM of AHA for 30 min (Click Chemistry Tools). At the end of the labeling period, bacterial cultures were treated with 100 μg/mL of chloramphenicol. Cells were collected by centrifugation at 4°C, washed thrice with ice-cold phosphate buffered saline (PBS) and stored at −80°C.

Cell pellets were thawed and re-suspended in a lysis buffer consisting of 50 mM Tris-HCl pH 8.0, 5% glycerol, 0.5% sodium dodecyl sulfate (SDS) and 1x protease inhibitor cocktail (Roche). Cells were lysed in a MiniBeadbeater-96 (BioSpec) and insoluble debris was removed by centrifugation (10 min, 10,000 X *g*, 4°C). Covalent attachment of fluorescent AFDye 488-alkyne (Click Chemistry Tools) to AHA containing proteins was carried out using Click-&-Go™ Protein Reaction Buffer Kit (Click Chemistry Tools) according to the manufacturer’s instructions. Protein concentrations were determined using a Pierce BCA Protein Assay Kit (Thermo Fisher Scientific). Fluorescent signals in samples were measured in a SpectraMax i3x plate reader (Molecular Devices) with 480 nm excitation and 520 nm emission wavelengths. The rate of protein synthesis was estimated as the fluorescence signal normalized by the protein content of the sample.

To determine if alterations on protein synthesis were systemic, AF488-labeled samples were separated by SDS-PAGE and fluorescence in gels was captured with an Amersham Imager 680 (GE Healthcare Life Sciences). To ensure that equal amounts of protein were loaded in each lane, gels were subsequently stained using the ProteoSilver™ Plus Silver Stain Kit (Sigma).

### MgtC immunoblot analysis

Wild-type (14028s) or *mgtC* (EL4) *Salmonella* were grown in MOPS medium containing 10 μM MgCl_2_ and 500 μM K_2_HPO_4_ for the indicated amount of time. Equivalent amounts of bacterial cells normalized by OD600 values were collected, washed with PBS, suspended in 0.15 mL SDS sample buffer (Laemmli sample buffer), and boiled. Cell extracts were loaded and resolved using 4–12% NuPAGE gels (Life Technologies). Proteins were then electro-transferred onto nitrocellulose membrane (iBlot; Life Technologies) following the manufacturer’s protocol. MgtC was detected using polyclonal anti-MgtC antibody (90) and the secondary antibody, horseradish peroxidase-conjugated anti-mouse IgG fragment (GE). The blots were developed with the SuperSignal West Femto Chemiluminescent system (Pierce) and visualized with an Amersham Imager 600 (GE Healthcare Life Sciences). Mouse-anti RpoB antibody (Thermo Fisher Scientific) was used as the loading control.

### Image acquisition, analysis and manipulation

Plates, gel and membrane images were acquired using an Amersham Imager 600 (GE Healthcare Life Sciences). ImageJ software (91) was used to crop the edges and adjust the brightness and contrast of the images. These modifications were simultaneously performed across the entire set images to be shown.

## Acknowledgments

M. H. P. is supported by grant AI148774 from the National Institutes of Health and funds from The Pennsylvania State University College of Medicine. E.A.G. is supported by grants AI49561 and AI120558 from the National Institutes of Health.

---

**Table S1.** Bacterial strains and plasmids used in this study

| Strain | Relevant characteristics | Source |
| --- | --- | --- |
| ***Escherichia coli*** |  |  |
| EC100D | *pir*<sup>+</sup> (DHFR) host strain used for generation and propagation of plasmid constructs | Epicentre |
| MG1655 | F<sup>-</sup> no $\gamma\delta$ λS (K12) | (92) |
| ***Salmonella enterica* serovar Typhimurium** |  |  |
| 14028s | wild-type | (93) |
| EL4 | Δ*mgtC* | (30) |
| MP24 | Δ*atpB*::Km<sup>R</sup> | (93) |
| MP25 | Δ*mgtC* Δ*atpB*::Km<sup>R</sup> | This study |
| MP1251 | Δ*pitA*::Ap<sup>R</sup> | This study |
| MP1252 | Δ*yjbB*::Km<sup>R</sup> | This study |
| MP1479p | Δ*pitA*::Ap<sup>R</sup> Δ*yjbB*::Km<sup>R</sup> | This study |
| MP1254 | Δ*mgtC* Δ*pitA*::Ap<sup>R</sup> | This study |
| MP1255 | Δ*mgtC* Δ*yjbB*::Km<sup>R</sup> | This study |
| MP1720 | Δ*mgtC* Δ*pstSCAB*::Km<sup>R</sup> | This study |
| RB39 | Δ*pitA* Δ*pstSCAB*::Km<sup>R</sup> Δ*yjbB* (3ΔPi) | This study |
| MP1665 | *phoR*<sup>L421A</sup> | This study |
| **Plasmids** |  |  |
| pSIM6 | rep<sub>pSC101</sub><sup>ts</sup> Amp<sup>R</sup> P<sub>CI857</sub>-γ βexo | (81) |
| pCP20 | rep<sub>pSC101</sub><sup>ts</sup> $\lambda$ cI857 FLP Amp<sup>R</sup> Cm<sup>R</sup> | (94) |
| pKD4 | rep<sub>R6Kγ</sub> Amp<sup>R</sup> FRT Km<sup>R</sup> FRT | (95) |
| pKD4-Ap<sup>r</sup> | rep<sub>R6Kγ</sub> Amp<sup>R</sup> FRT Apr<sup>R</sup> FRT | (96) |
| pSLC-242 | rep<sub>R6Kγ</sub> Amp<sup>R</sup> FRT Cm<sup>R</sup> *PrhaB-relE* FRT | (89) |
| pUHE-21-2-*lacI*<sup>q</sup> | rep<sub>pMB1</sub> *lacI*<sup>q</sup> Amp<sup>R</sup> vector control | (86) |
| pUHE-MgtC | rep<sub>pMB1</sub> *lacI*<sup>q</sup> Amp<sup>R</sup> Plac-*mgtC* | (97) |
| pUHE-PstSCAB | rep<sub>pMB1</sub> *lacI*<sup>q</sup> Amp<sup>R</sup> Plac-*pstSCAB* | This study |
| pUHE-PmrB | rep<sub>pMB1</sub> *lacI*<sup>q</sup> Amp<sup>R</sup> Plac-*pmrB* | (98) |
| pGFPc | rep<sub>p15A</sub> Cm<sup>R</sup> promoterless *gfp* vector control | (15) |
| pPstS-GFPc | rep<sub>p15A</sub> Cm<sup>R</sup> *PpstS-gfp* | (15) |
| pPmgtCB-GFPc | rep<sub>p15A</sub> Cm<sup>R</sup> *PmgtCB-gfp* | This study |
| pPphoB-GFPc | rep<sub>p15A</sub> Cm<sup>R</sup> *PphoB-gfp* | (15) |
| pGFP<sub>AAV</sub> | rep<sub>pMB1</sub> Amp<sup>R</sup> promoterless *gfp*<sub>AAV</sub> vector control | (15) |
| pPpstS-GFP<sub>AAV</sub> | rep<sub>pMB1</sub> Amp<sup>R</sup> *PpstS-gfp*<sub>AAV</sub> | (15) |
| pBbB2k-GFP | rep<sub>BBR1</sub> Km<sup>R</sup> *tetR* Ptet-*gfp* | (87) |
| pBbB2K-AtpAGD | rep<sub>BBR1</sub> Km<sup>R</sup> *tetR* Ptet-*atpAGD* | This study |

**Table S2.**
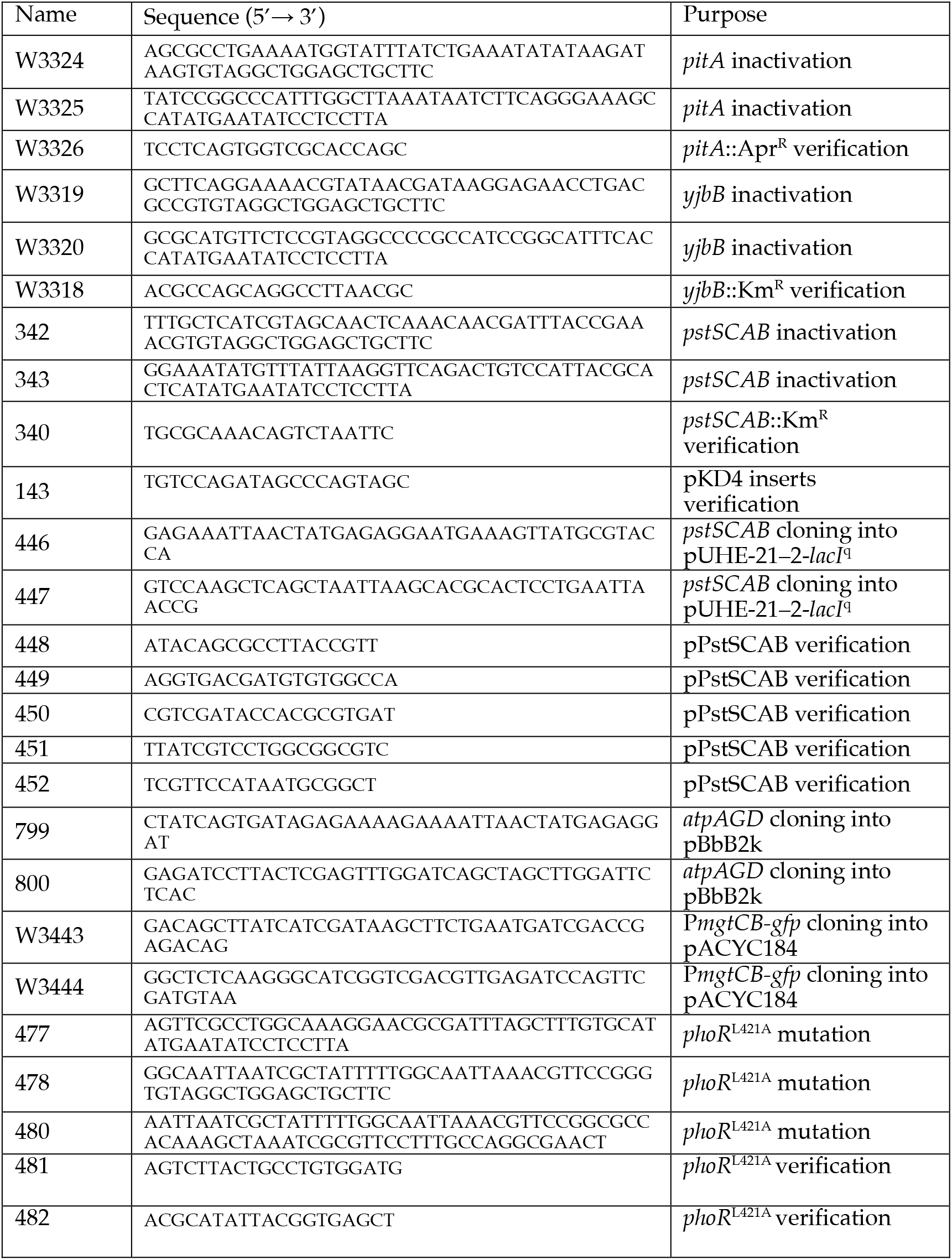
Oligonucleotides sequences used in this study

**Figure S1.**
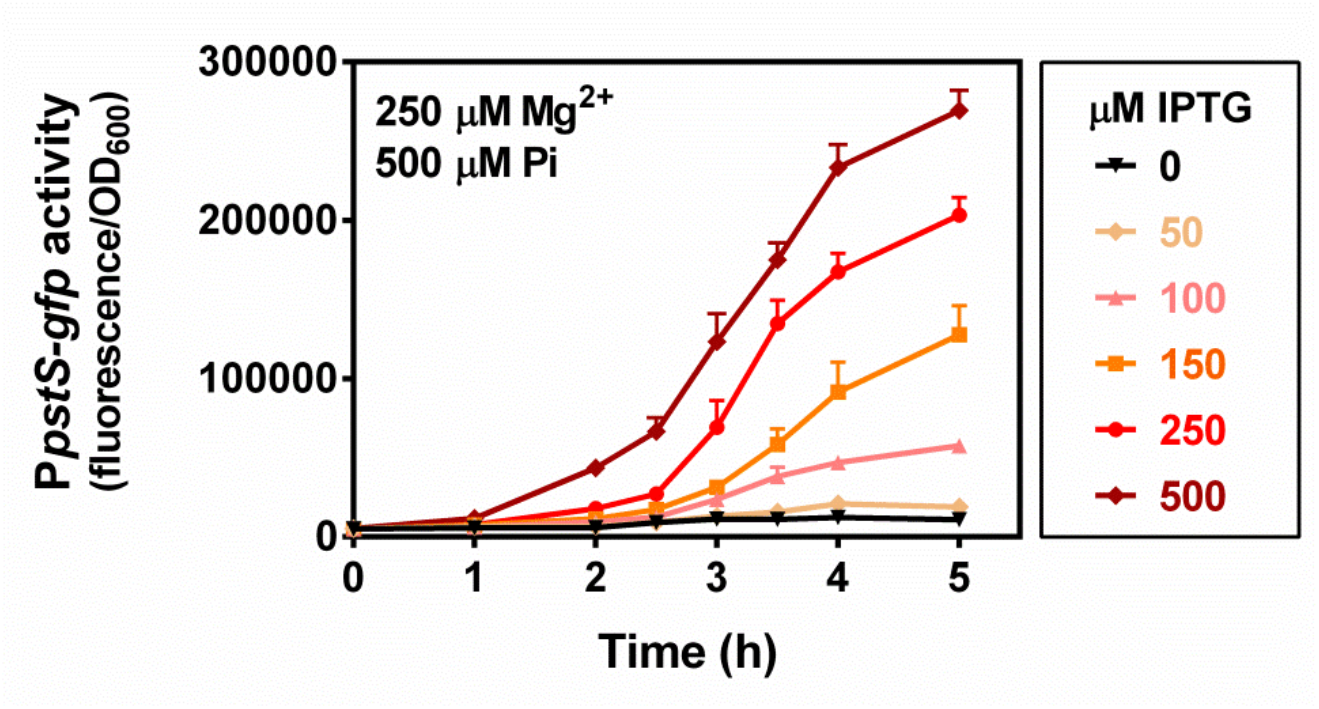
MgtC elicits a dose-dependent activation of the PhoB-activated PstSCAB transporter. Fluorescence from wild-type (14028s) *Salmonella* carrying pP*pstS*-GFPc during growth in MOPS medium containing 250 μM MgCl_2_, 500 μM K_2_HPO_4_, and the indicated IPTG concentrations. Means ± SDs of three independent experiments are shown.

**Figure S2.**
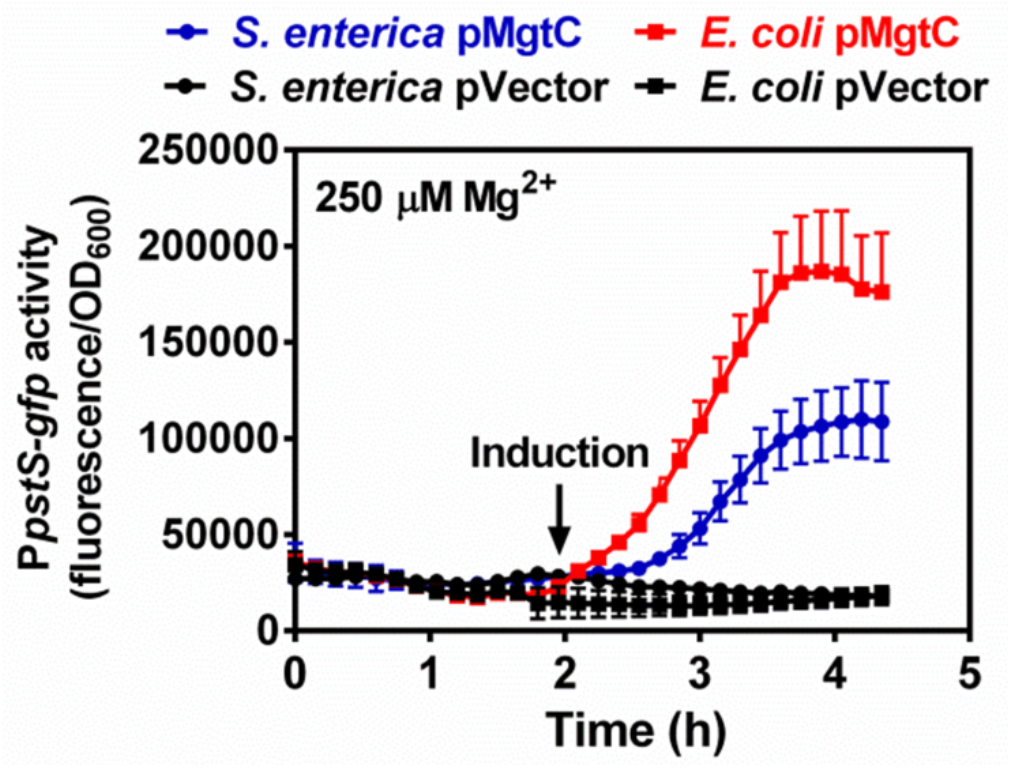
Heterologous expression of *Salmonella* MgtC in *E. coli* activates the PhoB/PhoR two-component system. Fluorescence from wild-type (14028s) *Salmonella enterica,* and wild-type *Escherichia coli* (MG1655) carrying *pPpstS-* GFPc and either pMgtC (pUHE-MgtC) or pVector (pUHE-21). Cells were grown in MOPS medium containing 250 μM MgCl_2_ and 500 μM K_2_HPO_4_. 250 μM IPTG was added to the cultures following 2 h of growth. Means ± SDs of three independent experiments are shown.

**Figure S3.**
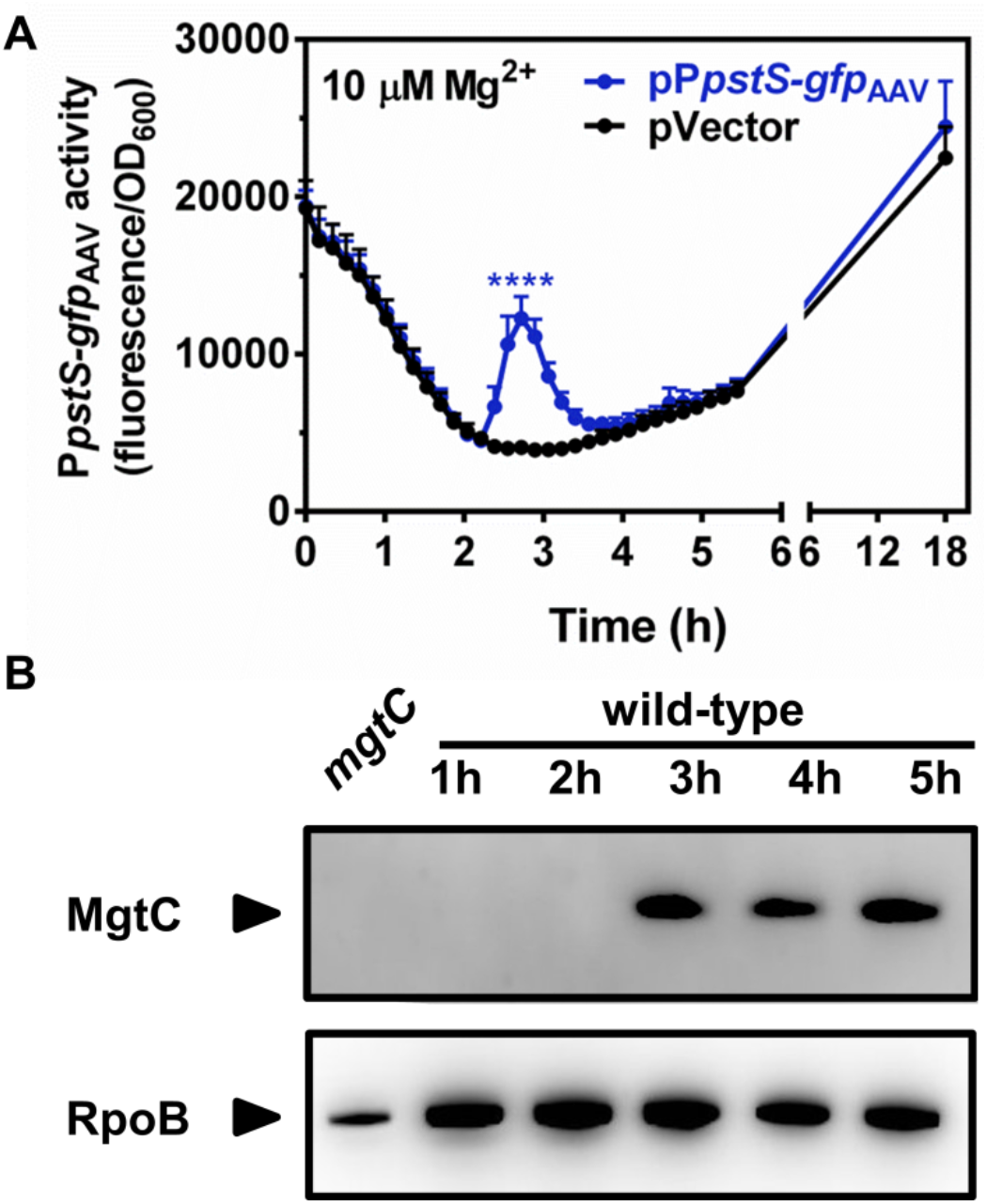
Expression timing of *pstSCAB* and MgtC during cytoplasmic Mg^2+^ starvation. *(A)* Fluorescence from wild-type (14028s) *Salmonella* carrying pP*pstS-*GFP_AAV_ or pVector (the promoterless GFP_AAV_ vector pGFP_AAV_) plasmid. Note that unstable GFP variants such as GFP_AAV_ enable monitoring of activation and silencing of gene expression (99). Cells were grown in MOPS medium containing 10 μM MgCl_2_ and 500 μM K_2_HPO_4_. Means ± SDs are shown. *(B)* Immunoblot analysis using anti-MgtC (upper panel) or anti-RpoB (lower panel, loading control) antibodies of crude extracts prepared from wild-type (14028s) or *mgtC* (EL4) *Salmonella* at the indicated timepoints. Similar expression timings are obtained when transcriptional fusions of *phoB-gfp* and *mgtC-gfp* are used instead (15). For A and B, cells were grown in MOPS medium containing 10 μM MgCl_2_ and 500 μM K_2_HPO_4_. Graphs and images are representative of three independent experiments.

**Figure S4.**
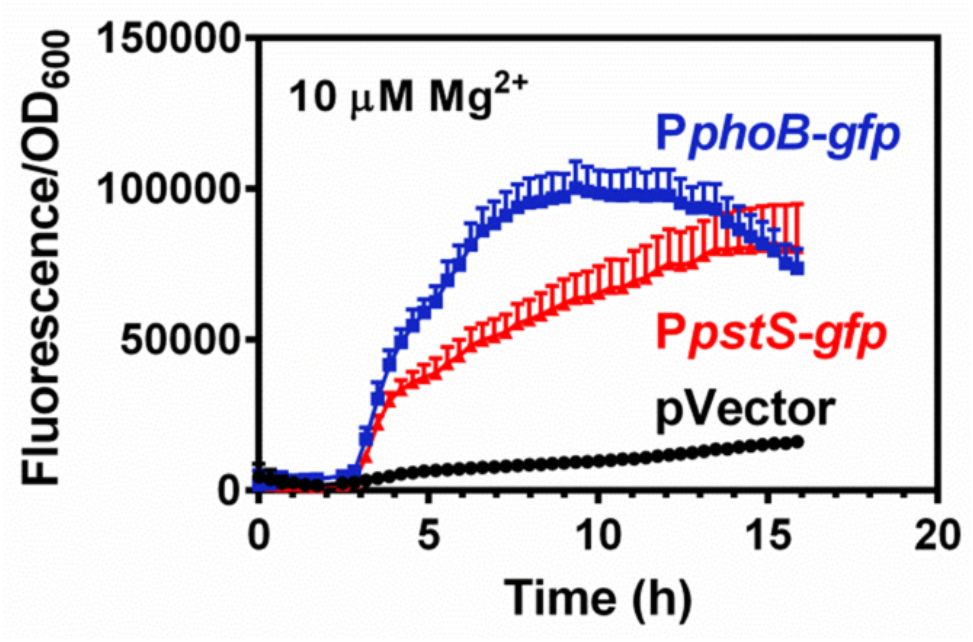
Effect of Mg^2+^ starvation on transcription of *phoB* and *pstSCAB* in *E. coli*. Fluorescence from wild-type (MG1655) *E. coli* carrying pP*phoB*-GFPc, pP*pstS-*GFPc, or pVector (the promoterless GFP plasmid pGFPc). Cells were grown in MOPS medium containing 10 μM MgCl_2_ and 500 μM K_2_HPO_4_. Means ± SDs of three independent experiments are shown.

**Figure S5.**
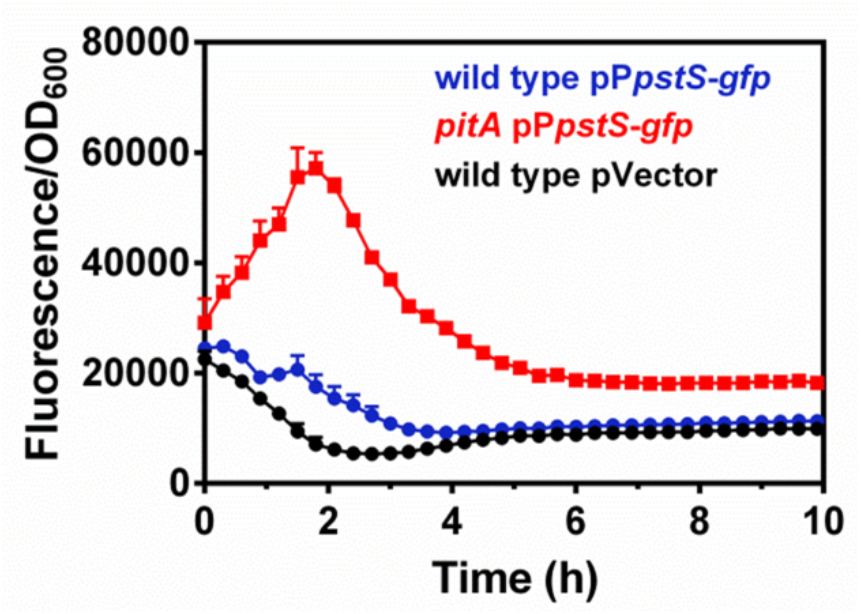
Effect of deleting *pitA* Pi transporter on *pstS* transcription. Fluorescence from wild-type (14028s) or *pitA* (MP1251) *Salmonella* carrying pP*pstS-*GFPc or pVector (the promoterless GFP plasmid pGFPc). Cells were grown in MOPS medium containing 250 μM MgCl_2_ and 1000 μM K_2_HPO_4_. Means ± SDs of three independent experiments are shown.

**Fig. S6.**
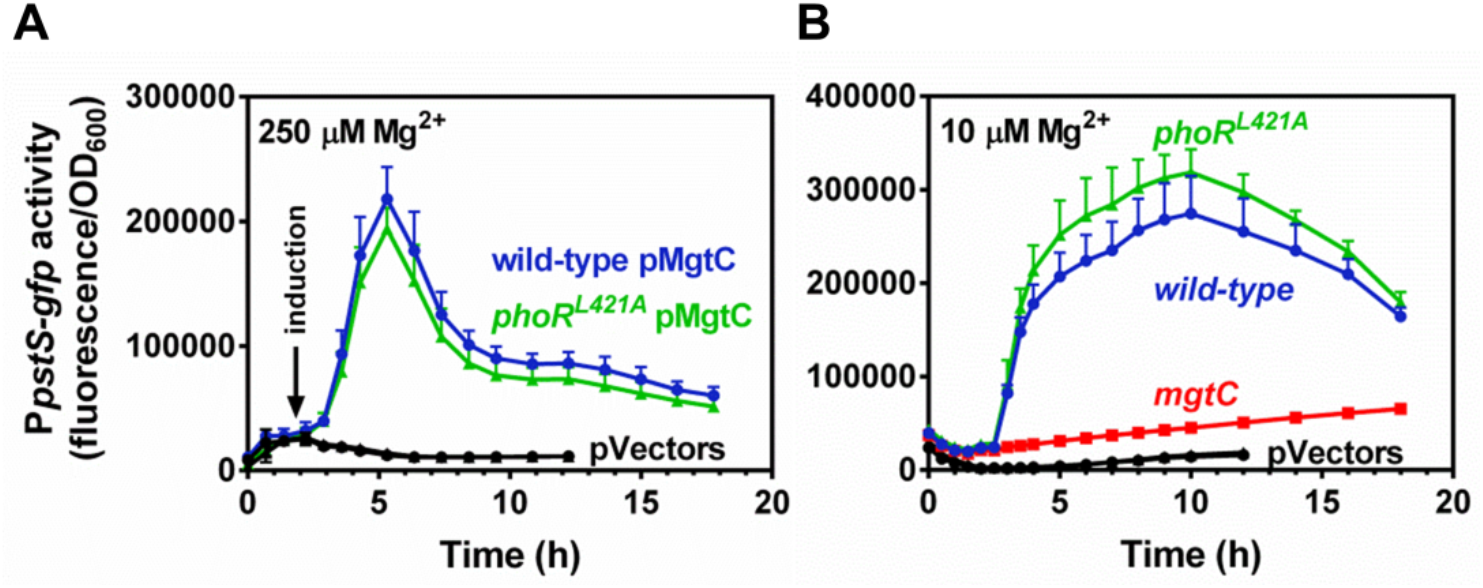
Effect of *phoR*^L421A^ substitution on the activation of the PhoB/PhoR two-component system. *(A)* Fluorescence from wild-type (14028s) and *phoPy*^L421A^ (MP1665) *Salmonella* carrying pP*psfS*-GFPc and either pMgtC (pUHE-MgtC) or pVector (pUHE-21). Cells were grown in MOPS containing 250 μM MgCl_2_ and 500 μM K_2_HPO_4_. Following 2 h of growth, 250 μM IPTG was added to the cultures. *(B)* Fluorescence from wild-type (14028s) and *phoR*^L421A^ (MP1665) *Salmonella* carrying pPstS-GFPc or pVector (the promoterless GFP plasmid pGFPc). Cells were grown in MOPS containing 10 μM MgCl_2_ and 500 μM K_2_HPO_4_. Means ± SDs of at least three independent experiments are shown.

